# Aberrant Transitional Alveolar Epithelial Cells Promote Pathogenic Activation of Lung Fibroblasts in Preclinical Fibrosis Modeling

**DOI:** 10.1101/2024.06.17.599351

**Authors:** Evan T Hoffman, Willy Roque Barboza, Luis R Rodriguez, Rachna Dherwani, Yaniv Tomer, Aditi Murthy, Adam Bennett, Ana Nottingham, Apoorva Babu, Katrina Chavez, Charlotte H Cooper, Maria C Basil, Micha Sam Brickman Raredon, Jeremy Katzen

## Abstract

Pulmonary fibrosis (PF) is a chronic progressive lung disease histopathologically characterized by fibrotic remodeling and the presence of pathological epithelial and mesenchymal cell populations in the distal lung parenchyma. Within the epithelial compartment, a subset of alveolar type 2 cells (AT2s) enter and persist in an aberrant transitional state. Whether and how these aberrant transitional cells participate in lung fibrosis is not known. To address this, we exploited the *Sftpc^C121G^* mouse model, where we previously demonstrated that chronic expression of a PF-associated point mutation (*C121G*) in the AT2-specific surfactant protein C (*Sftpc*) gene results in spontaneous and progressive fibrosis driven by intrinsic AT2 dysfunction. We utilized single cell RNA sequencing to demonstrate the emergence of pathologic epithelial and mesenchymal cells in the *Sftpc^C121G^* murine lung fibrosis model, including aberrant transitional alveolar epithelial cells as well as transitional and fibrotic fibroblasts. Aberrant transitional alveolar epithelial cells share similar transcriptional profiles to human aberrant basaloid cells, including the upregulation of profibrotic gene markers (*Fn1, Ctgf, Tgfb2, Pdgfb, Spp1*), and develop a unique interactome with pathogenic lung fibroblasts. We developed a method to reliably flow sort aberrant transitional alveolar epithelial cells, and we highlight their ability to cause fibrotic activation of fibroblasts in *ex vivo* organoid assays and using conditioned supernatant, suggesting a profibrotic secretome. We conclude that aberrant transitional alveolar epithelial cells actively contribute to fibrotic lung remodeling through pathogenic activation of alveolar fibroblasts.

## Introduction

Pulmonary Fibrosis (PF) is the most common adult interstitial lung disease, with rising incidence and prevalence (1, 2) and a clinical course marked by progressive loss of lung function resulting in lung transplant or death within 3-7 years of diagnosis (3). Despite the substantial and growing public health burden of PF, there are only two FDA approved medications, which only modestly slow lung function decline (4, 5). A major barrier to developing better PF therapies is the failure to model and study the proximal cellular dysfunction that initiates disease (6).

A dominant hypothesis in the field is that intrinsic AT2 dysfunction is the proximal pathogenic event in PF (3, 7). AT2s have a high protein biosynthetic burden and high metabolic requirements, making them prone to intrinsic and extrinsic insults that initiate stress pathways (7). Among the specific signaling pathways identified in aberrant AT2s in both patients with advanced (8, 9) and “early/preclinical” (10) PF are those involved in regulating protein homeostasis (“proteostasis”) (10, 11), suggesting that this specific stress is a proximal cellular event in PF pathogenesis. Supporting this, a subset of PF patients develop disease from mutations in the BRICHOS domain of the AT2-restricted surfactant protein C (*SFTPC*) gene (12–14) that generate a misfolded aggregation-prone proprotein (proSP-C) that induces proteostatic stress (15, 16) (17). Leveraging these mutations, we developed and published the *Sftpc^C121G^* model of spontaneous lung fibrosis from knock-in of a *Sftpc* BRICHOS mutation (C121G) in the adult mouse lung, allowing us to study intrinsic AT2 dysfunction in lung fibrosis (18–20). However, a direct link between a dysfunctional AT2 and pathogenic fibroblasts, the primary effector cells of fibrotic lung remodeling, has not been established.

Defining histopathologic features of human PF and murine fibrotic lungs include anomalous alveolar epithelial cells and an extracellular matrix-producing fibroblast population. We and multiple other groups have shown through murine modeling that, following severe fibrotic lung injury, AT2s enter a transitional state defined by the expression of specific cytokeratins (*Krt8*, *Krt19*) and *Cldn4* (referred to as PATS, DAPTs, or Krt8+ cells), and that a subset of these cells become “stuck”, failing to fully differentiate to AT1s (19–28). However, functional characterization of the disease-associated epithelial and fibroblast cell states has been limited by the difficulty of isolating them to interrogate their interactions. We overcame this barrier in our *Sftpc^C121G^* model by identifying a cell-surface receptor that is uniquely expressed on the aberrant transitional AT2s during lung fibrosis, which allowed us to isolate these cells. Using this capacity, we tested the hypothesis that aberrant transitional AT2s enriched in profibrotic mediators can directly promote fibrogenic activation in alveolar fibroblasts. We show that the secreted factors from this pathogenic subset of AT2s indeed elicits fibrotic activation in fibroblasts, and in doing so we demonstrate the previously suggested but unestablished profibrotic role of the lung epithelium in pulmonary fibrosis.

## Results

### Identification of an aberrant profibrotic alveolar transitional cell subset in an *Sftpc^C121G^* model of pulmonary fibrosis

As previously reported, *Sftpc^C121G^* mice receiving a weekly dose of oral gavage TMX (200mg/kg) developed spontaneous and progressive lung fibrosis without preceding alveolitis or the need for a secondary extrinsic lung injury (**Figure 1A,B**) (19). We sought to further characterize the epithelial and mesenchymal cell populations in *Sftpc^C121G^* mice at the end-stage of the model as a means of understanding fibrotic signaling pathways that may contribute to pulmonary fibrosis. To accomplish this, we performed single cell RNA sequencing (scRNAseq) at week 7 of the model on whole lung epithelial (EpCAM^+^) and mesenchymal (EpCAM^-^;CD31^-^;CD45^-^) cells from two *Sftpc^C121G^* and two *Sftpc^WT^* mice (**Figure 1A,C**). After quality filtering, we obtained 29,303 cell transcriptomic profiles that, based on marker gene expression, showed representation from the lung mesenchyme (*Pdgfra, Pdgfrb*, *Acta2*), proximal epithelium (*EpCam, Krt5, Mcgb1a1, Muc5b, Foxj1*), and distal epithelium (*EpCam, Sftpc, Abca3, Hopx, Ager*) (**Figure 1C, Supplemental Figure 1**). Similar to previous scRNAseq studies on end-stage human PF patient lung samples, we observed significant transcriptomic alterations in all three cell compartments as evidenced by shifts in clustering between *Sftpc^WT^* and *Sftpc^C121G^* samples (**Figure 1C, Supplemental Figure 1**) (21, 23).

**Figure 1.**
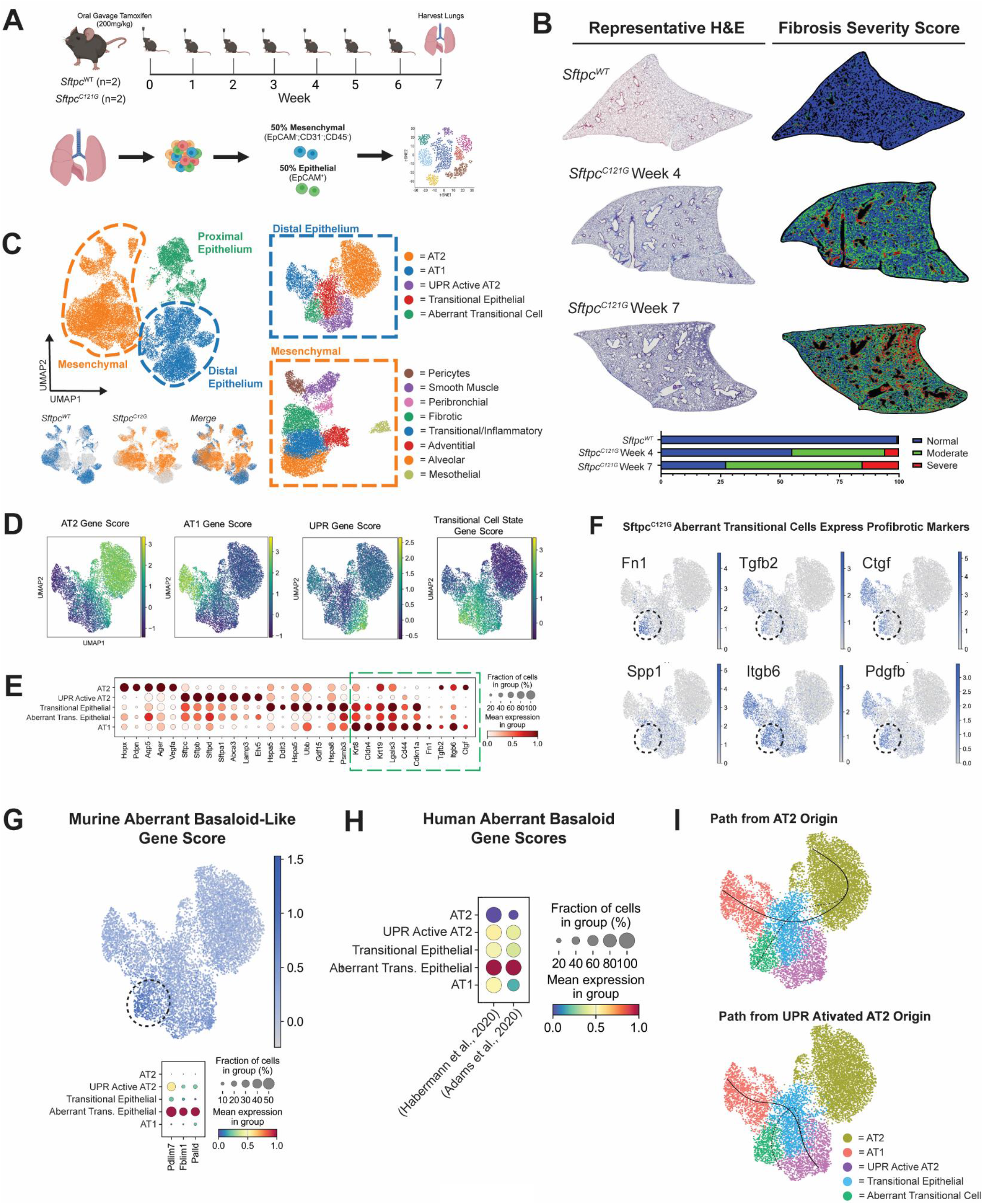
Identification of key pathologic epithelial and mesenchymal cell types in the chronic *Sftpc^C121G^* mutation model. (A) Schematic of chronic tamoxifen-induced Sftpc^C121G^ mutation model and subsequent cell sorting for single-cell RNA sequencing (scRNA-seq). (B) Representative H&E staining with fibrosis severity scoring (normal = blue, moderate = green, and severe = red) and bar graph showing relative amounts of injury per lung lobe. (C) scRNA-seq UMAP representation of mesenchymal, proximal (airway), and distal (alveolar) epithelial subsets in both Sftpc*^WT^* and Sftpc*^C121G^* mice. Inserts depict annotated subset analysis of distal epithelial and mesenchymal populations. UPR, unfolded protein response. (D) UMAP representation of gene marker scoring of distal epithelial populations. (E) Dot plot of individual gene expression in distal epithelial cell populations. (F) UMAP representation of defined profibrotic genes enriched within the aberrant transitional cell population. (G) Gene score representation by UMAP and dot plots of signature gene markers characterized in murine basaloid-like cells (26). (H) Gene score dot plots from human aberrant basaloid populations (21, 23). (I) Trajectory analysis suggesting aberrant transitional cells are a terminal cell fate in the *Sftpc^C121G^* alveolar epithelium.

To further characterize the alveolar epithelial cell populations, we isolated and re-clustered the distal lung epithelium from both *Sftpc^WT^* and *Sftpc^C121G^* mice (**Figure 1D-F, Supplemental Figure 2**). We observed that the *Sftpc^C121G^* alveolar epithelium had significantly reduced expression of homeostatic AT2 and AT1 markers, as previously described in human PF (**Figure 1D,E, Supplemental Figure 2**) (23). In contrast, we found the alveolar epithelium in *Sftpc^C121G^* mice primarily upregulated genes associated with UPR activation (*Hspa5, DDit3, Gdf15*) and an alveolar epithelial transitional cell state (*Krt8, Krt19*, *Lgals3*) (18–25). We additionally identified a specific cluster that we termed “aberrant transitional cells” based on high expression of transitional cell markers (*Krt8, Cldn4, Krt19*) as well as profibrotic (*Fn1, Ctgf, Tgfb2, Spp1*), and senescence marker *Cdkn1a* (**Figure 1E,F**). We found that this cell population closely resembled the transcriptomic profile of a previously described “basaloid” like subset of mouse transitional cells (**Figure 1G**), including the expression of defining gene markers *Pdlim7, Fblim1,* and *Palld,* and shared a similar transcriptomic profile to aberrant basaloid cells described in human PF (**Figure 1H, Supplemental Table 2**) (26–28). Trajectory inference analysis starting from either the homeostatic AT2 cluster or the *Sftpc^C121G^* UPR-enriched cluster demonstrated two trajectories—one that the marked the AT2-to-AT1 progression seen in functional epithelial repair and a second that terminated in the aberrant transitional cell cluster suggesting this is a terminal cell fate (**Figure 1I**). Together this analysis of the *Sftpc^C121G^* alveolar epithelium revealed the presence of an aberrant transitional cell in a stalled differentiation state that shares transcriptomic features of the human PF aberrant basaloid cells.

### Chronic *Sftpc^C121G^* lungs possess mesenchymal cell populations found in PF with alterations in epithelial-mesenchymal crosstalk

We next sought to characterize distinct mesenchymal populations that were present at end stage fibrosis in the *Sftpc^C121G^* lungs based on known heterogeneity of lung fibroblasts within healthy adult murine lungs, as well as shifts in gene expression and cell states within fibrotic lung tissues (24, 29–33). Mesenchymal cell clusters were identified as expressing fibroblast markers (*Pdgfra, Pdgfrb*, *Acta2*), and excluding epithelial markers (*EpCAM*) (**Figure 1C, Supplemental Figure 1B**). Cluster analysis from the concatenated *Sftpc^WT^* and *Sftpc^C121G^* lungs revealed 8 mesenchymal cell populations with distinct gene profiles (**Supplemental Figure 3**). This analysis included clusters of alveolar (*Col13a1, Npnt, Inmt*) and adventitial (*Col14a1, Pi16, Ly6a*) fibroblasts found primarily in *Sftpc^WT^* lungs, as well as fibrotic (*Cthrc1, Col1a1, Tgfb1, Fst, Spp1)* and a recently described transitional (*Lcn2, Hp, Sfrp1*) fibroblast population that were derived almost exclusively from *Sftpc^C121G^* lungs (**Supplemental Figure 3**) (29, 30, 32, 33).

To investigate aberrant signaling that may occur between epithelial and mesenchymal cell populations in *Sftpc^C121G^* fibrotic lungs, we utilized a recently developed software package, NICHES, that allowed for unbiased ligand-receptor mapping at a single cell resolution (34). As NICHES accounts for directionality of signaling, we were able to analyze both epithelial-to-mesenchymal (**Figure 2**) and mesenchymal-to-epithelial (**Supplemental Figure 4**) signaling. With regards to epithelial-to-mesenchymal signaling, we discovered that alveolar epithelial and mesenchymal cells segregated into 11 clusters based on ligand-receptor interactions (**Figure 2A,B**). Samples derived from *Sftpc^WT^* and *Sftpc^C121G^* mice displayed minimal clustering overlap, suggesting that a divergent interactome develops in the fibrotic model (**Figure 2A**), with Clusters 0,1,4,5 comprised almost exclusively of *Sftpc^C121G^* cells (**Figure 2B**). In particular, the sending population in Cluster 5 was largely comprised of aberrant transitional cells (84.0%) and the receiving mesenchymal populations were fibrotic (32.0%) and transitional (22.5%) fibroblasts (**Figure 2C**). Further analysis of Cluster 5 showed it was defined by established profibrotic ligand-receptor signaling pairs, including *Tgfb2-Tgfbr2*, *Pdgfb-Pdgfrb, Ctgf-Lrp1,* and *Spp1-Cd44* (**Figure 2D,E**) (35–40), suggesting a potential role of aberrant transitional cells in promoting a fibrotic phenotype in lung mesenchymal cells through direct signals.

**Figure 2.**
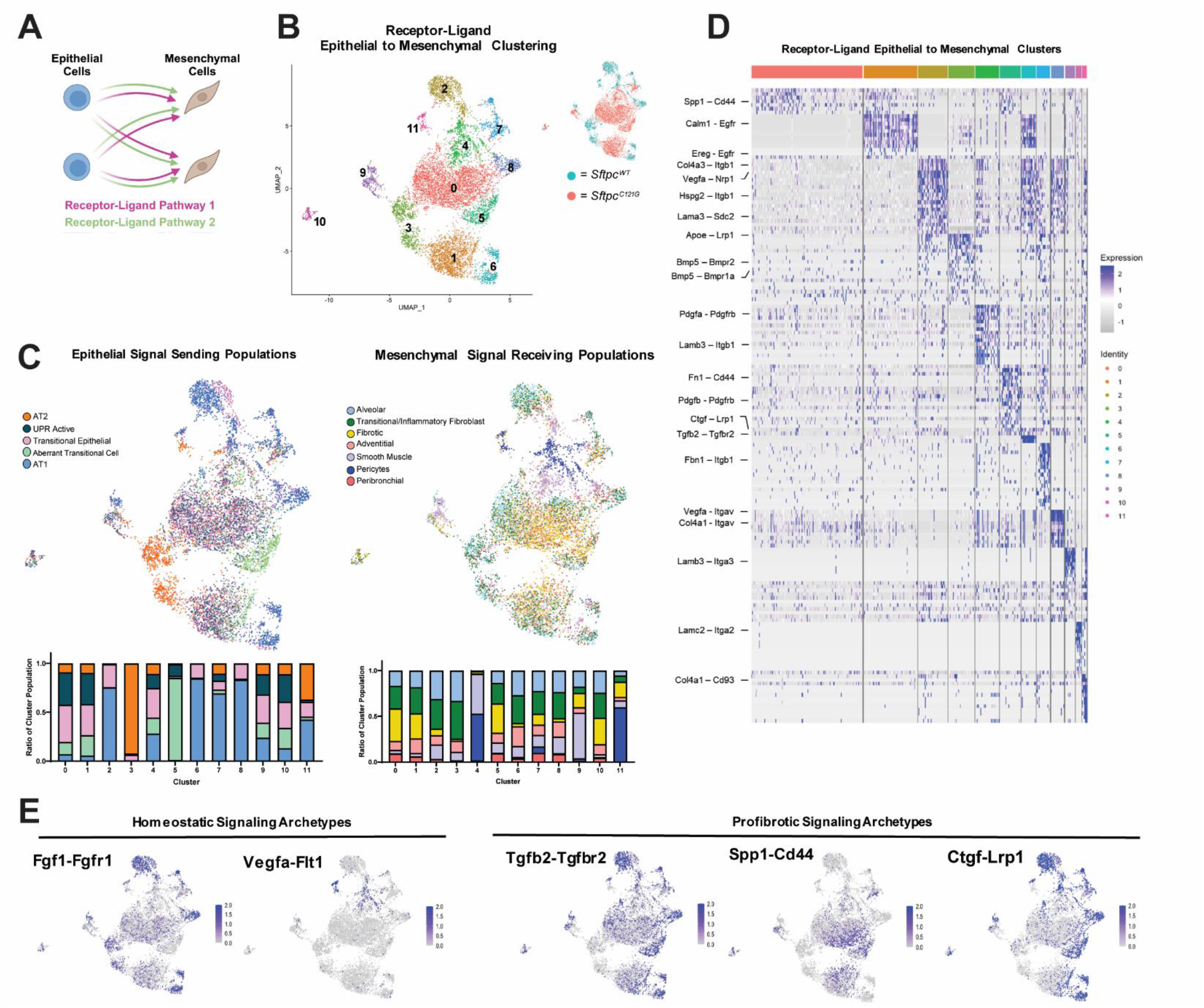
Epithelial-to-mesenchymal signaling is characterized by a loss of homeostatic signaling and gain of profibrotic signaling. (A) Schematic demonstrating single-cell ligand-receptor mapping using NICHES. (B) UMAP representation of unbiased epithelial-to-mesenchymal ligand-receptor mapping clustered dependent on distinct signaling archetypes. (C) UMAPs labeled by epithelial sending cell type and mesenchymal receiving cell type populations. Ratios of each cell type within a given cluster is provided. (D) Heatmap of individual ligand-receptor pairs that define each cluster archetype. (E) Representative UMAPs of known homeostatic and profibrotic ligand-receptor pairs that are altered in fibrosis.

NICHES analysis of the mesenchymal-to-epithelial signaling also demonstrated aberrant signaling in *Sftpc^C121G^* lungs (**Supplemental Figure 4**). Re-clustering cell populations according to mesenchymal-to-epithelial ligand-receptor signaling revealed 8 separate clusters, with Clusters 2,3,8 primarily associated with *Sftpc^WT^* cells and Clusters 0,1,4,5 primarily associated with *Sftpc^C121G^* cells. Aberrant transitional cells clustered with fibrotic and transitional fibroblasts (Clusters 0, 4, and 5) **(Supplemental Figure 4A-C**) suggesting they received ligand signals from these pathogenic fibroblast populations. These clusters were defined by enrichment of ligand-receptor pairs indicative of dysregulated TGF-β signaling and increased extracellular matrix interactions (*Col1a1-Itgb3, Col1a1-Itga5, Tgfb1-Itgav,* and *Fn1-Cd44*) **(Supplemental Figure 4C,D**). We also discovered a loss of homeostatic signaling cues in *Sftpc^C121G^* lungs, including decreased *Fgf7-Fgfr2* and *Bmp4-Bmpr1a* signaling and increased Fst*- Bmpr2* signaling, suggesting impaired alveolar niche maintenance in *Sftpc^C121G^* lungs **(Supplemental Figure 4E**) (31, 41). Together, the ligand-receptor mapping analysis revealed that the aberrant transitional cells were implicated in bidirectional fibrogenic signaling with pathological fibroblasts.

### Flow sorted aberrant transitional cells retain profibrotic gene expression

While our scRNAseq analysis of *Sftpc^C121G^* lungs and studies performed by others in both human PF and mouse fibrosis models (21, 23, 26–28), describe a similar subset of transitional cells that express profibrotic markers, the isolation and *ex vivo* analysis of these cell populations has remained elusive (42, 43). Our scRNAseq analysis revealed that *Sftpc^C121G^* aberrant transitional cells were enriched in expression of *Cd44*, which transcribes the transmembrane glycoprotein CD44 (**Figure 3A**). Confirming that these cells locate to regions of fibrotic remodeling, immunofluorescent staining of lung sections demonstrated that KRT8+/CD44+ transitional cells were found within regions of dense collagen deposition in *Sftpc^C121G^* peripheral lung tissue (**Figure 3B; Supplemental Figure 5B,C**). Based on this, we developed a sorting strategy for isolating CD44+/- alveolar epithelial cells by FACS (**Figure 3C; Supplemental Figure 5A**). We observe that 8.39% (± 1.74 SD) of AT2s in *Sftpc^WT^* lungs are positive for CD44, similar to previous studies that showed these cells in the homeostatic lung have unique stem cell properties (44). In *Sftpc^C121G^* lungs, we found a significant increase in CD44+ alveolar epithelial cells to 40.25% (± 4.20 SD) (**Figure 3D**). Bulk RNA sequencing from flow sorted CD44+/- alveolar epithelial cells from *Sftpc^WT^* and *Sftpc^C121G^* lungs (referred to herein as CD44+/- *Sftpc^WT^* AT2s or CD44+/- *Sftpc^C121G^* AT2s, respectively) provided evidence of CD44 as a useful marker for isolating aberrant transitional cell from *Sftpc^C121G^* lungs (**Figure 3E-H**). We found that CD44+ *Sftpc^C121G^* AT2s are enriched in transitional (*Krt19, Cldn4*) and profibrotic *(Fn1, Tgfb2, Spp1, Pdgfb*) genes (**Figure 3F,G**), and are enriched in extracellular matrix organization by pathway analysis (**Figure 3H**). Further, we found that both CD44+ and CD44- *Sftpc^C121G^* AT2s downregulated *Fgfr2* expression, suggesting a reduced capacity to respond to homeostatic alveolar niche signaling (**Figure 3F**) (31). Additionally, CD44+ *Sftpc^WT^* AT2s had increased gene expression of inflammatory (*Il-1*β*, Tnf, Ccl20*), proliferation (Mki67), and AT1 (*Spock2,* Rtkn2) markers, corroborating a recent finding that CD44+ *Sftpc^WT^* AT2s are a proinflammatory population of AT2s with increased stem-like properties (**Figure 3F**) (45).

**Figure 3.**
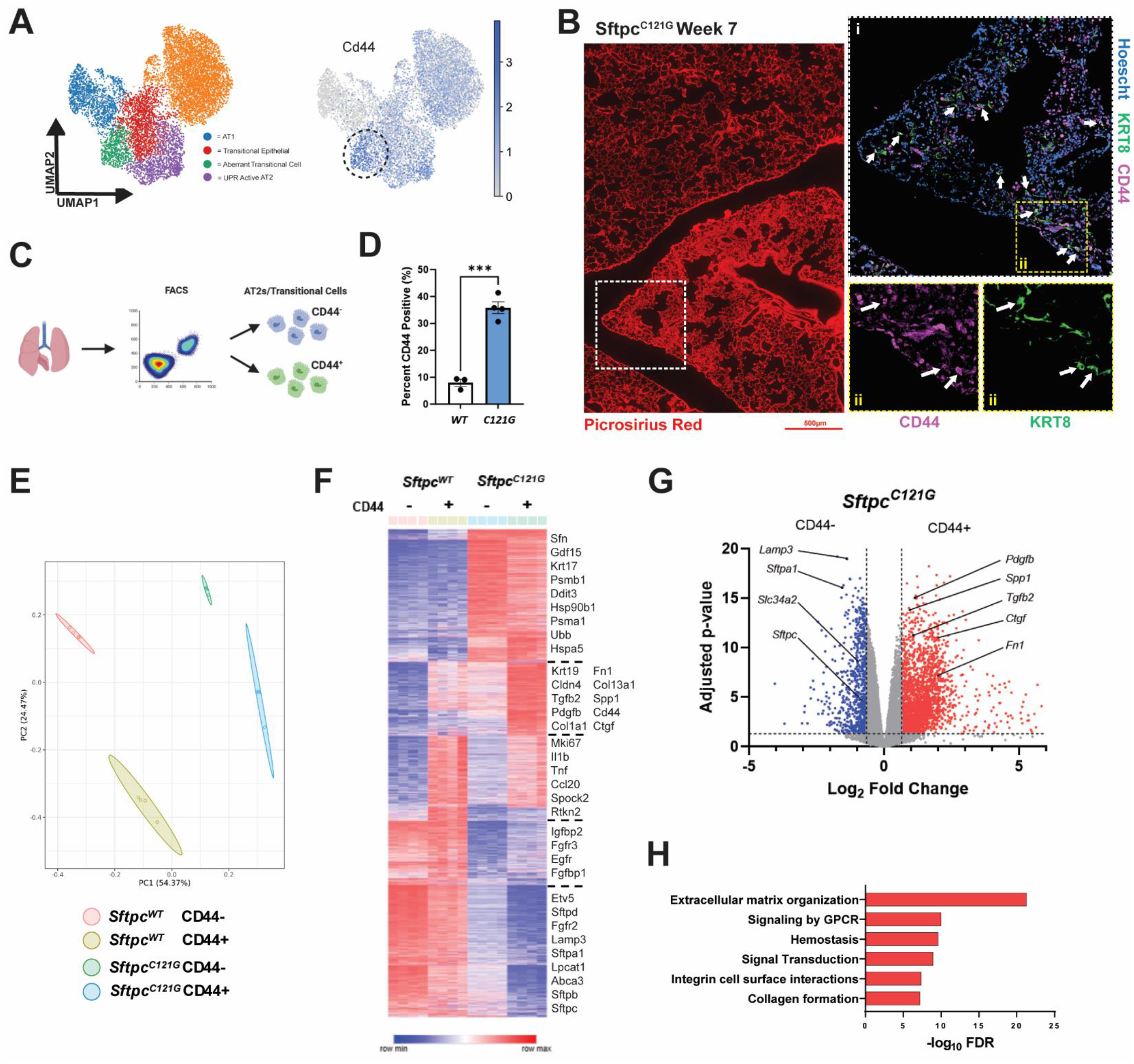
Sorted aberrant CD44+ transitional cells are enriched in profibrotic mediators. (A) UMAP representation of *Cd44* gene enrichment in aberrant transitional cells. (B) Immunofluorescent chemistry in chronic Sftpc*^C121G^* mice lungs showing CD44+/Krt8+ transitional cells proximal in areas of high collagen content by PSR. (C) Schematic for flow sorting CD44+/- AT2/transitional cells. (D) Percent CD44+/- AT2/transitional cells in *Sftpc^WT^* and *Sftpc^C121G^* mice identified by flow cytometry. (E) PCA demonstrating bulk transcriptomic difference between CD44+/- AT2 and transitional cells from *Sftpc^WT^* and *Sftpc^C121G^* mice. (F) Heatmap of top DEGs within each condition. (G) Volcano plot showing differentially upregulated (red) and downregulated (blue) genes (>1.5 fold change, adjusted p value <0.05) between CD44- and CD44+ *Sftpc^C121G^* AT2s/ transitional cells. (G) Gene ontogeny analysis for upregulated biological processes in CD44+ *Sftpc^C121G^*, compared to CD44- *Sftpc^C121G^*, AT2s and transitional cells.

To determine whether *Cd44* expression in aberrant transitional cells was unique to the *Sftpc^C121G^* model, we analyzed CD44+ cells from a separate preclinical model of PF. Utilizing a previously published scRNAseq dataset (24), we found that a subset of alveolar transitional cells in the single-dose bleomycin mouse model also had increased expression of *Cd44* (**Supplemental Figure 6A**). These *Cd44^hi^* transitional cells emerged during the fibrogenic phase (days 9-14) of the bleomycin model and expressed similar profibrotic genes as CD44+ *Sftpc^C121G^* AT2s, most notably *Fn1* (**Supplemental Figure 6B,C**). To determine whether CD44+ AT2s are present in the model, we administered bleomycin to lineage traceable *Sftpc^CreERT2^;Rosa26^tdT^* mice and harvested lungs during the fibrotic phase, at day 14 post-bleomycin (**Supplemental Figure 6D**). In fibrotic lungs, we observed elongated tdT+/CD44+ cells by IFC as well as a significant increase in tdT+/CD44+ cells by flow cytometry—32.60% (± 3.32 SD) compared to 3.57% (± 2.32 SD) in healthy lungs (**Supplemental Figure 6E,F**). Additionally, flow sorted tdT+/CD44+ cells from fibrotic lungs had increased expression of transitional cell (*Cldn4*) and profibrotic markers (*Fn1, Tgfb2*), suggesting that CD44 is a useful marker for isolating aberrant transitional cells in various models of pulmonary fibrosis (**Supplemental Figure 6G**). We were thus able to develop a sorting strategy to isolate a distinct subset of alveolar epithelial cells that emerge during fibrosis modeling and that are enriched in profibrotic signaling ligands.

### CD44+ *Sftpc^C121G^* AT2s contribute to profibrotic epithelial-mesenchymal crosstalk

While previous studies and our transcriptomic analysis suggest that transitional alveolar epithelial cells may contribute to pulmonary fibrosis by signaling to fibroblasts, the direct impact of these cells and how they respond to reciprocal signals from fibroblasts has yet to be described (21, 23, 26–28). To address this, we interrogated CD44+ *Sftpc^C121G^* AT2s through *ex vivo* co-culture modeling. We developed an *ex vivo* mixed culture organoid assay in which *Sftpc^WT^* AT2s or CD44+/- *Sftpc^C121G^* AT2s were co-cultured with either *Sftpc^WT^* or *Sftpc^C121G^* fibrotic lung fibroblasts (**Figure 4A**). Comparison of sorted bulk Pdgfra+ fibroblasts (Pdgfra^+^;CD45^-^;EpCAM^-^;CD31^-^;MCAM^-^) from *Sftpc^WT^* or Week 7 *Sftpc^C121G^* fibrotic lungs (referred to herein as *Sftpc^WT^* or *Sftpc^C121G^* fibroblasts, respectively) demonstrated *Sftpc^C121G^* fibroblasts had increased expression of both transitional (*Lcn2, Sfrp1, Hp*) and fibrotic (*Cthrc1, Col1a1*) marker genes (**Supplemental Figure 7A,B**). Upon mixed organoid culture, we found that *Sftpc^C121G^* fibroblasts significantly increased colony forming efficiency (CFE) and size of *Sftpc^WT^* AT2 organoids, and size of CD44- *Sftpc^C121G^* AT2 organoids, while no difference in CFE or size were observed in CD44+ *Sftpc^C121G^* AT2 organoids (**Figure 4B-D**). These data demonstrate that *Sftpc^C121G^* fibroblasts can drive AT2 proliferation, but CD44+ *Sftpc^C121G^* AT2 lacked this response. This suggests the response to mesenchymal cell signaling on AT2 proliferation may be dependent on AT2 cell state.

**Figure 4.**
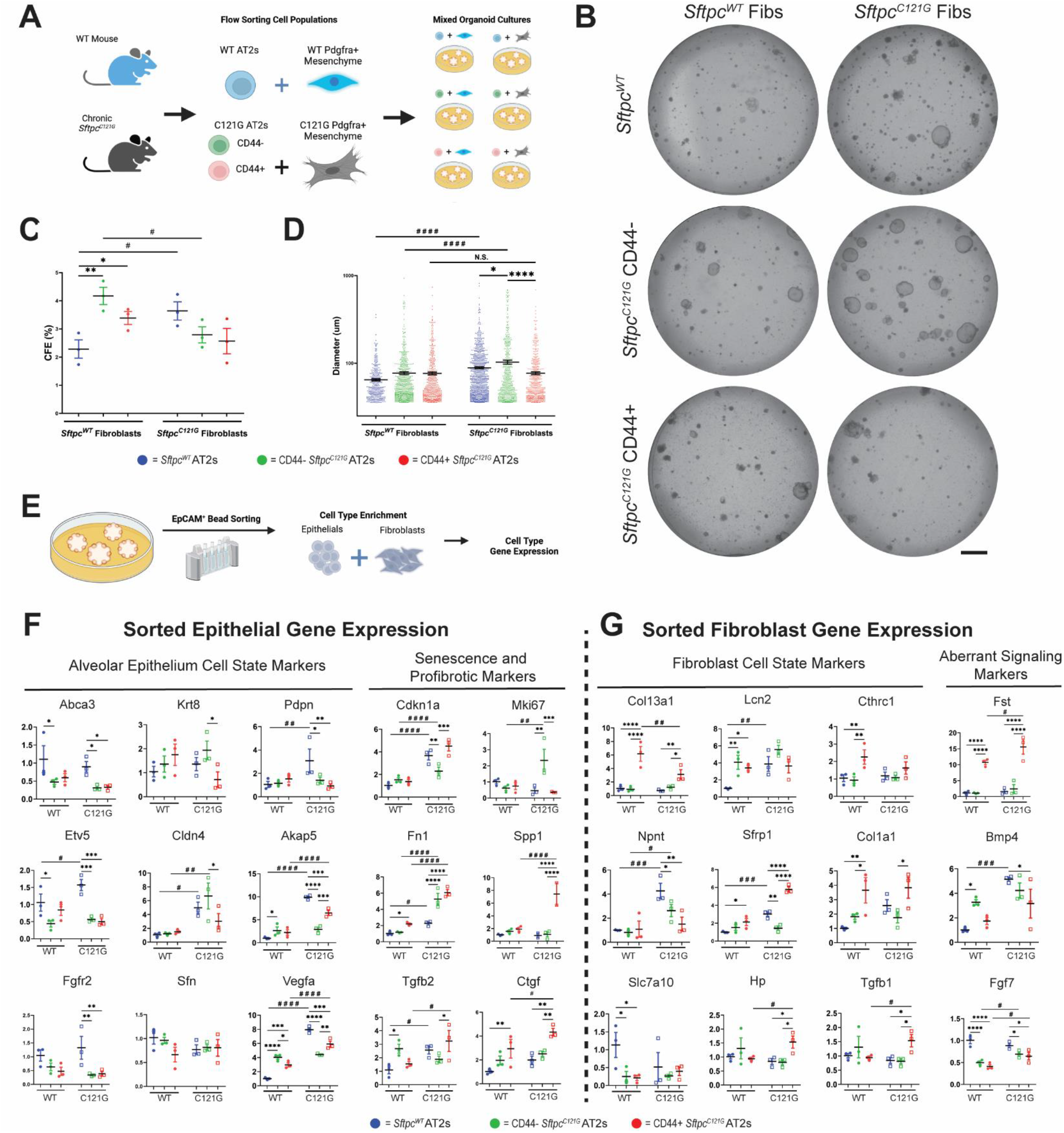
Mixed organoid culture demonstrates CD44+ *Sftpc^C121G^* AT2s induce a pathologic response in fibroblasts. (A) Schematic depicting mixed organoid cultures derived from various combinations of AT2/transitional cells and bulk Pdgfra^+^ mesenchymal cells from *Sftpc^WT^* and *Sftpc^C121G^* mice. (B) Representative imaging of organoid cultures. Scale bar = 1mm. (C) Colony forming efficiency (CFE; %) and (D) organoid diameter (µm) and of organoids at Day 14 in culture. (E) Schematic depicting EpCAM+ magnetic bead sorting from Day 14 organoids prior to gene expression analysis. (F) Gene expression analysis from enriched epithelial or fibroblast populations derived from Day 14 organoids. Ordinary one-way ANOVA with uncorrected Fisher’s LSD multiple comparisons were performed. (*) were used when comparing samples cultured within the same fibroblast condition. (#) were used when comparing samples within the same epithelial condition. */# P<0.5, **/## P<0.01, ***/### P<0.001, ****/#### P<0.0001.

To determine the signaling mechanisms which may drive differences seen in mixed organoid cultures, we developed an isolation assay in which EpCAM+ magnetic bead sorting was used to isolate cells from organoid cultures (**Figure 4E**), allowing us to reliably distinguish both epithelial-enriched and fibroblast-enriched cell populations for gene expression analysis (**Supplemental Figure 7C**). As expected, based on our transcriptomic data, AT2 markers (*Abca3, Etv5, Fgfr2*) were downregulated in both CD44+ and CD44- *Sftpc^C121G^* AT2s compared to *Sftpc^WT^* AT2s, regardless of fibroblast origin in co-culture (**Figure 4F**). Strikingly, compared to *Sftpc^WT^* AT2s, we observed decreased expression in AT1 markers (*Pdpn, Akap5, Vegfa*) in both CD44+ and CD44- *Sftpc^C121G^* AT2s when cultured with *Sftpc^C121G^* fibroblasts, suggesting impaired AT2 to AT1 differentiation. Conversely, we observed increased expression of *Akap5* and *Vegfa* when cultured with *Sftpc^WT^* fibroblasts, suggesting a potential role for homeostatic lung fibroblasts in rescuing impaired *Sftpc^C121G^* alveolar epithelial cell differentiation (**Figure 4F**). Further, CD44+ and CD44- *Sftpc^C121G^* AT2s had increased expression of profibrotic markers (*Fn1, Spp1, Tgfb2, Ctgf*), particularly when cultured with *Sftpc^C121G^* fibroblasts, as well as increased *Cdkn1a* expression, suggesting a senescent phenotype in CD44+ *Sftpc^C121G^* epithelial cells *ex vivo* (**Figure 4F**). These data reveal a differential effect of fibrotic lung fibroblasts on homeostatic lung (*Sftpc^WT^*) AT2s and CD44+ *Sftpc^C121G^* epithelial cells, with a potential role in amplifying the aberrant transitional cell phenotype.

Based upon the increase in profibrotic genes in co-cultured CD44+ and CD44- *Sftpc^C121G^* AT2s we next sought to answer whether these cells induce a profibrotic response in co-cultured fibroblasts. We identified evidence that CD44+ *Sftpc^C121G^* AT2s indeed drove a fibrotic phenotype in fibroblasts (**Figure 4G**). CD44+ *Sftpc^C121G^* AT2s cultured with *Sftpc^WT^* fibroblasts resulted in increased fibroblast expression of transitional (*Sfrp1*) and fibrotic (*Cthrc1, Col1a1, Fst*) marker genes. CD44+ *Sftpc^C121G^* AT2s cultured with fibrotic *Sftpc^C121G^* fibroblasts resulted in decreased expression of homeostatic alveolar fibroblast marker *Npnt* and further increased transitional (*Sfrp1, Hp*) and fibrotic (*Cthrc1, Col1a1, Tgfb1*, *Fst*) marker genes. We thus found that in organoid co-culture CD44+ *Sftpc^C121G^* AT2s could directly impact the alveolar fibroblast phenotype, driving expression of pathogenic fibroblast marker genes in *Sftpc^WT^* fibroblasts and amplifying the pathologic phenotype of fibrotic *Sftpc^C121G^* lung fibroblasts.

### Altered FGF7 signaling contributes to impaired transitional alveolar epithelial cell progenitor capacity

Having identified loss of homeostatic niche signaling archetypes between pathogenic fibroblasts and CD44+ *Sftpc^C121G^* AT2s (**Supplemental Figure 4E**), we sought to connect this with the progenitor cell defect we observed when these populations were placed in organoid co-culture (**Figure 4A-D**). We observed that both CD44+ and CD44- *Sftpc^C121G^* AT2s appeared to inhibit *Fgf7* expression in both *Sftpc^WT^* and *Sftpc^C121G^* fibroblasts, and that CD44+ *Sftpc^C121G^* AT2s decreased *Bmp4* expression in *Sftpc^C121G^* fibroblasts (**Figure 4G**), suggesting that aberrant transitional cells may alter homeostatic alveolar niche signaling pathways (31, 41). Given the decreased expression of *Fgfr2* in CD44+ and CD44-*Sftpc^C121G^* AT2s (**Figure 3F**), and their capacity to decrease *Fgf7* expression in both *Sftpc^WT^* and *Sftpc^C121G^* fibroblasts (**Figure 4F**), we asked if CD44+ and CD44- *Sftpc^C121G^* AT2s could respond to homeostatic FGF7 signaling. To address this question, we utilized a previously published AT2-alone organoid model, in which AT2s may be cultured independently from feeder cells (46, 47). Using this model with an FGF7 dose response (**Figure 5A**), we observed increasing colony forming efficiency (CFE) in *Sftpc^WT^* AT2s with each increase in FGF7 concentration (4, 10, and 25ng/mL), while both CD44+ and CD44- *Sftpc^C121G^* AT2s exhibited no change in CFE (**Figure 5B,C**). However, we did observe increased organoid diameter in CD44+ and CD44- *Sftpc^C121G^* AT2s with increasing FGF7 concentration, suggesting a potential subset of these cells is capable of responding to FGF7. Interestingly, we also observed increased AT1 gene marker expression (*Pdpn, Aqp5*) in *Sftpc^WT^* AT2s at the highest FGF7 concentration compared to CD44+ and CD44- *Sftpc^C121G^* AT2s, further suggesting impaired AT1 differentiation potential in *Sftpc^C121G^* alveolar epithelial cells (**Figure 5D**). These data suggest that transitional alveolar epithelial cells may persist in the fibrotic lung by altered epithelial-mesenchymal crosstalk through FGF7 signaling.

**Figure 5.**
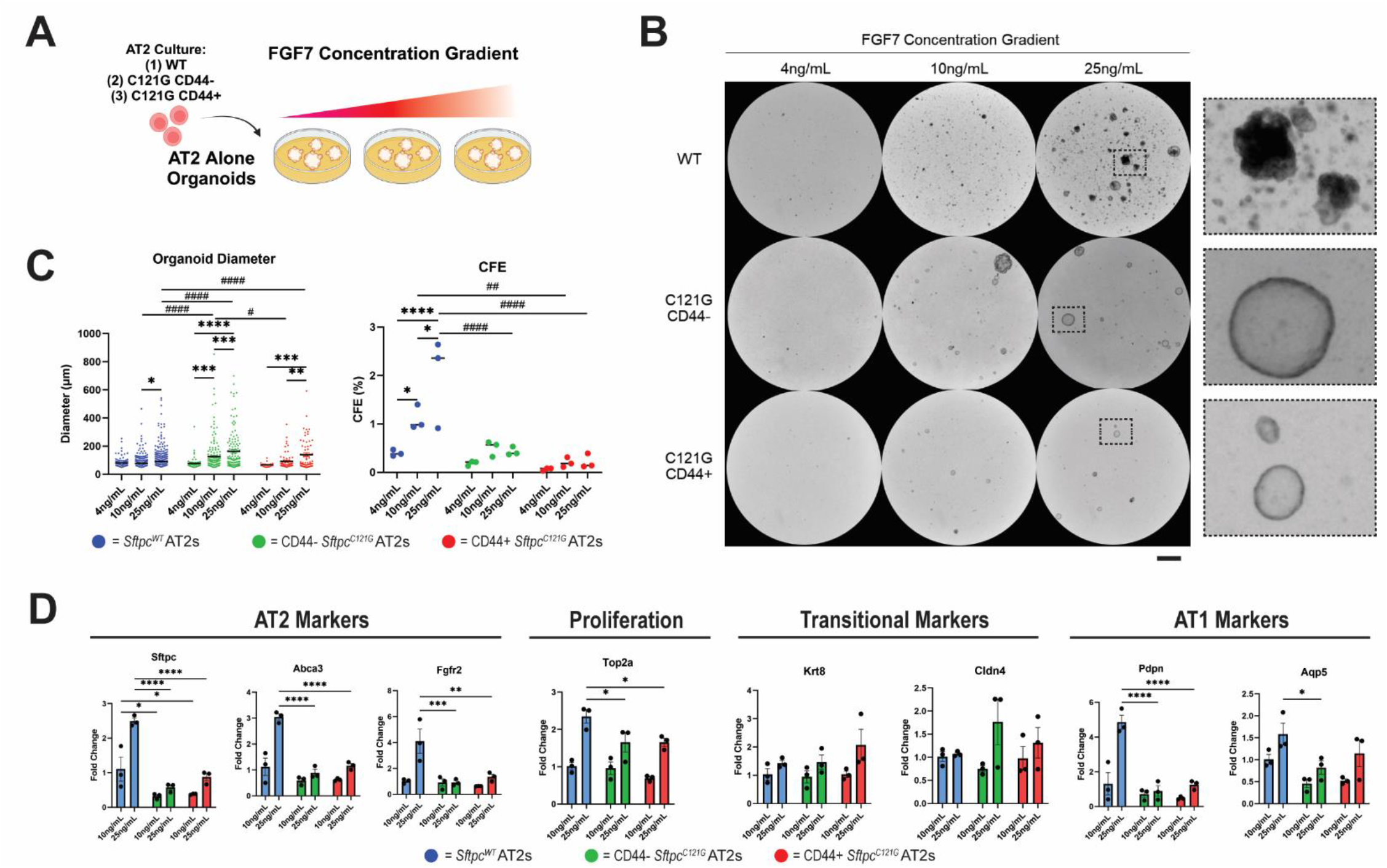
CD44+ *Sftpc^C121G^* AT2s do not proliferate in response to increasing FGF7 dosage. (A) Schematic depicting AT2-alone 3D culture with a gradient of FGF7 dosing. (B) Representative imaging of organoid cultures. (C) Organoid diameter (µm) and colony forming efficiency (CFE; %) of organoids at Day 14 in culture. Two-way ANOVA (* = significant difference between samples with the same origin, # = significant difference between samples with the same FGF7 dosing), */# P<0.5, **/## P<0.01, ***/### P<0.001, ****/#### P<0.0001. (D) Gene expression analysis of select gene markers of cultured cells. Two-way ANOVA, * P<0.5, ** P<0.01, *** P<0.001, **** P<0.0001.

### Supernatant from CD44+ *Sftpc^C121Gs^* drive a fibrotic phenotype in alveolar fibroblasts

Since mixed culture organoids suggested CD44+ *Sftpc^C121G^* AT2s are capable of driving fibroblasts towards a fibrotic phenotype, we wanted to determine whether this process was driven by soluble factors. Accordingly, *Sftpc^WT^* AT2s and CD44+/- *Sftpc^C121G^* AT2s were cultured in 2D Transwells and supernatant was collected (**Figure 6A**). Classically, this 2D assay has been used to assess AT2-to-AT1 differentiation, and we observed in all conditions a decrease in AT2 gene expression at Day 5 in culture, as expected (**Figure 6B**) (27, 48, 49). However, we found that CD44+ *Sftpc^C121G^* AT2s had reduced expression of AT1 markers (*Pdpn, Hopx*) at Day 5 compared to *Sftpc^WT^* AT2s, further suggesting impaired AT1 differentiation potential. While 2D culture led to an increase in profibrotic markers (*Tgfb2, Ctgf, Fn1*) in all alveolar epithelial cell conditions, CD44+ and CD44- *Sftpc^C121G^* AT2s expressed increased levels of these gene markers at Day 5 compared to *Sftpc^WT^* AT2s, and CD44+ *Sftpc^C121G^* AT2s expressed increased levels of *Ctgf* compared to CD44- *Sftpc^C121G^* AT2s.

**Figure 6.**
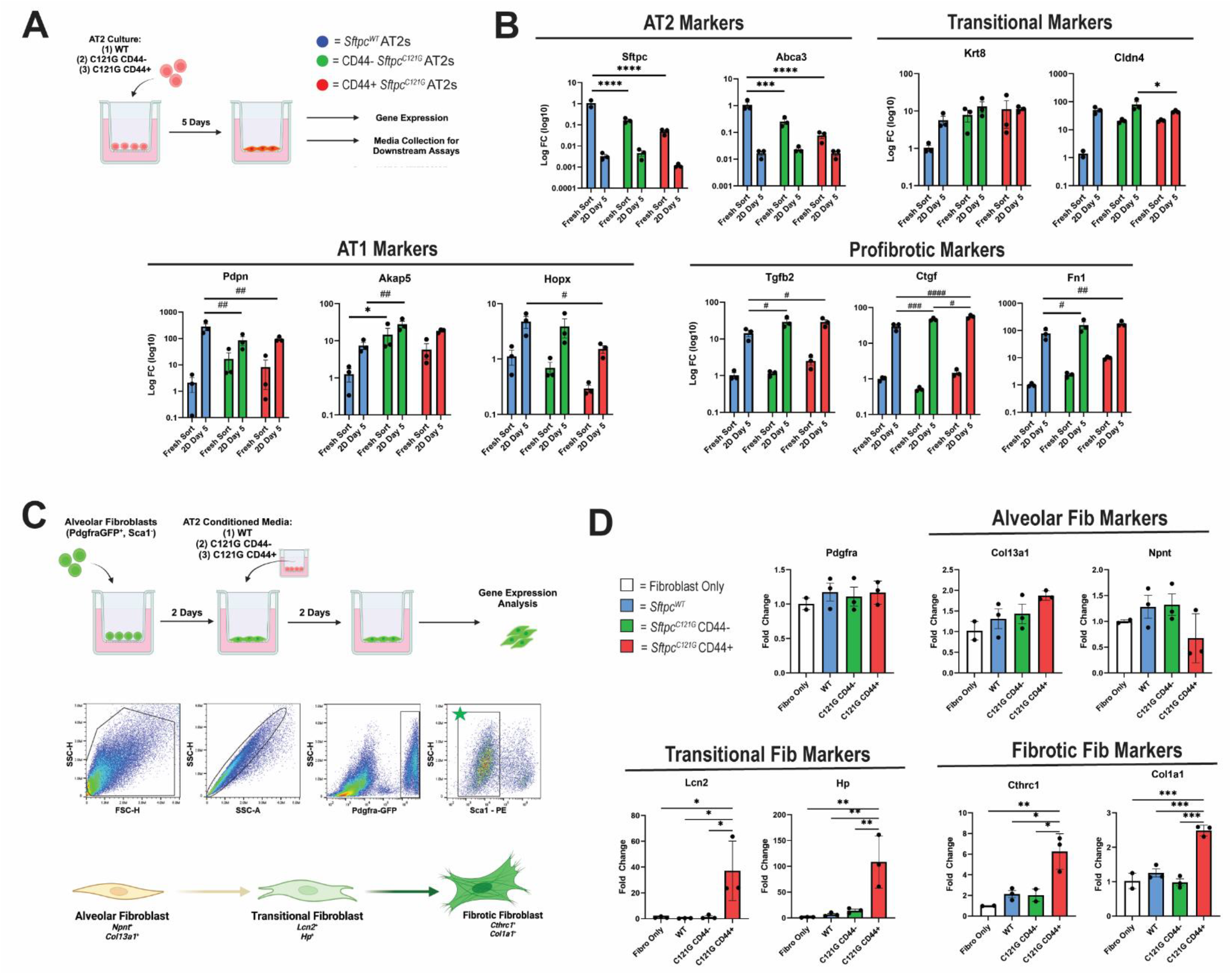
CD44+ *Sftpc^C121G^* transitional cells have impaired AT1 differentiation capacity and secrete soluble profibrotic mediators. (A) Schematic depicting gene expression analysis and supernatant collection from 2D cultures of *Sftpc^WT^* and CD44+/- *Sftpc^C121G^* AT2/transitional cells. (B) Gene expression of freshly sorted and 2D cultured cells at Day 5 in culture. Ordinary one-way ANOVA with uncorrected Fisher’s LSD multiple comparisons were performed. (*) were used when comparing freshly sorted samples. (#) were used when comparing 2D cultured samples. */# P<0.5, **/## P<0.01, ***/### P<0.001, ****/#### P<0.0001. (C) Schematic of healthy alveolar fibroblast exposure to conditioned supernatant from AT2s 2D cultured for five days including flow sorting strategy and cell state markers of alveolar, transitional, and fibrotic fibroblasts. (D) Gene expression of alveolar fibroblasts after two days of supernatant exposure. Statistical analysis by one-way ANOVA (*p < 0.05, ** p<0.005, *** p<0.0005).

At Day 5, supernatant from all alveolar epithelial cell cultures was collected and used as conditioned media for exposure studies on sorted alveolar fibroblasts (Pdgfra^+^;Sca1^-^) in 2D culture (**Figure 6C**). We observed significant upregulation of transitional (*Lcn2, Hp*) and fibrotic (*Cthrc1, Col1a1*) marker genes in alveolar fibroblasts exposed to supernatant from CD44+ *Sftpc^C121G^* AT2s, as compared to fibroblasts in basal media or fibroblasts treated with *Sftpc^WT^* AT2s or CD44- *Sftpc^C121G^* AT2 supernatant (**Figure 6D**). To confirm this finding was not a phenomenon specific to the chronic *Sftpc^C121G^* mouse model, we performed this parallel experiment using conditioned media from alveolar epithelial cells derived from bleomycin treated lungs, and observed that conditioned media from CD44+ AT2/transitional cell culture similarly elicited increased expression of *Lcn2, Hp,* and *Col1a1* in alveolar fibroblasts (**Supplemental Figure 6H,I**). We thus discovered that soluble factors from CD44+ AT2s from the fibrotic mouse lung, which develop a comparable gene signature to aberrant basaloid cells seen in human PF lungs, are capable of inducing a profibrotic response in alveolar fibroblasts, driving them into a pathogenic phenotype.

## Discussion

Pulmonary fibrosis (PF) is an ongoing clinical challenge for which the development of manageable and effective treatments is still necessary. Significant obstacles in therapy development include the identification of individual or multi-combinatorial signaling pathways that lead to ineffectual lung regeneration and the emergence of activated fibrotic fibroblast effector cells (50). In particular, aberrant transitional alveolar epithelial cells that emerge and accumulate in human PF and preclinical models of pulmonary fibrosis have been implicated in promoting fibrosis based on their transcriptomic profile and spatial localization (21–28). However, due to a lack of effective methods of isolating transitional epithelial cells, characterization of the profibrotic potential of these cells has relied primarily on genomic approaches. Here we report the identification of a subset of profibrotic aberrant transitional cells at the end stage of an *Sftpc^C121G^* preclinical mouse model of PF (**Figure 1**). We characterize, by scRNAseq and trajectory analysis, similarities between this population and aberrant basaloid cells observed in human IPF, including expression of profibrotic gene markers (*Fn1, Tgfb2, Ctgf, Spp1, Pdgfb*) and display transcriptomic characteristics of a stalled differentiation state, including upregulation of senescence-associated marker *Cdkn1a*. Additionally, we provide evidence, using bulk RNA sequencing, that the transmembrane glycoprotein, CD44, is an effective marker for flow sorting aberrant transitional cells for *ex vivo* modeling (**Figure 3**). Using this method, we isolate CD44+ *Sftpc^C121G^* AT2s and provide evidence of their bioactive role in promoting a fibrotic gene signature in alveolar fibroblasts in both organoid co-cultures and through secreted supernatant exposure assays (**Figures 4 and 6**). Together our findings illustrate that aberrant transitional alveolar epithelial cells secrete one, or multiple, factors that drive the emergence of fibrotic fibroblasts and appear to possess impaired differentiation capacity towards mature AT1s, suggesting their inability to regenerate injured alveolar epithelium. Further, flow sorting and *ex vivo* modeling strategies described in this study may serve as a platform for asking future questions about dysregulated signaling that occurs in PF as well as future therapeutic intervention studies.

In these studies we primarily relied on a chronic *Sftpc^C121G^* mutation mouse model, which we have previously demonstrated recapitulates the clinical course of PF (19). This fibrosis model leverages a clinically relevant mutation in the BRICHOS domain of the *Sftpc* gene, one of a class of over 60 mutations associated with PF, resulting in intrinsic AT2 ER stress and subsequent lung fibrosis (12–20). Importantly, chronic *Sftpc^C121G^* expression produces spontaneous and progressive end stage pathology without the need for extrinsic lung injury (i.e. bleomycin) or antecedent lung inflammation (19). We further validate this model through scRNAseq of end stage fibrotic lungs and demonstrate the emergence of PF-associated epithelial and mesenchymal cell populations (**Figure 1; Supplemental Figure 1-3**) (21, 23, 29, 30, 32). The ability to identify critical effector cells of PF, including transitional epithelial and mesenchymal populations (42, 43), highlights the utility of the chronic *Sftpc^C121G^* model for studying both early and end stage events that drive PF.

A major goal of this study was to elucidate epithelial-mesenchymal crosstalk signaling in PF, and to accomplish this we employed an innovative software package, NICHES, that allows for ligand-receptor mapping at a single cell level (34, 51). We were thus able to demonstrate signaling pathways that were altered in aberrant cell populations, most notably between aberrant transitional alveolar epithelial cells and transitional or fibrotic fibroblasts (**Figure 2; Supplemental Figure 4**). This included increased epithelial-to-mesenchymal interactions of multiple known profibrotic mediators (*Tgfb2-Tgfbr2, Ctgf-Lrp1, Pdgfb*-*Pdgfrb* and *Spp1-Cd44*) (35, 36). Importantly, unbiased ligand-receptor mapping allowed us to screen isolated CD44+ *Sftpc^C121Gs^* in *ex vivo* culture assays and confirm upregulation of profibrotic gene expression in both organoid (**Figure 4**) and conditioned media assays (**Figure 6**), which paralleled the activation of fibrotic fibroblasts. Notably, the increase in several profibrotic ligands within the *Sftpc^C121G^* alveolar epithelium further suggests the potential need for combinatorial therapeutics to reduce the activation of transitional and fibrotic fibroblasts (33, 50).

In addition to the expression of profibrotic ligands, we demonstrate that aberrant transitional alveolar epithelial cells appear to reside in a stalled differentiation state, likely in part due to aberrant bidirectional epithelial-mesenchymal crosstalk. Similar to previous studies in both human basaloid cells in PF (21, 52) and murine basaloid-like alveolar transitional cells (26), we show using trajectory analysis, dysregulated AT2-to-AT1 differentiation in aberrant transitional cells in *Sftpc^C121G^* mice (**Figure 1I**). Further, *ex vivo* modeling of *Sftpc^C121G^* alveolar epithelial cells appeared to provide additional evidence for impaired AT1 differentiation at gene expression level. Using a 2D culture method classically acknowledged to drive AT2-to-AT1 differentiation, we observed decreased expression of AT1 markers in CD44+ *Sftpc^C121G^* AT2s compared to *Sftpc^WT^* AT2s (**Figure 6**). This correlated with increased expression of profibrotic markers, including *Tgfb2*, which has previously been described to induce cell cycle arrest and inhibit AT2 differentiation *in vitro*, and suggests the potential for autocrine TGF-β signaling in impaired differentiation (27). Similarly, in an *ex vivo* organoid co-culture model, CD44+ and CD44- *Sftpc^C121G^* AT2s cultured with *Sftpc^C121G^* fibroblasts had decreased AT1 marker expression, compared to *Sftpc^WT^* AT2s, while CD44+ *Sftpc^C121G^* AT2s further expressed senescence maker, *Cdkn1a*, and profibrotic markers *Fn1* and *Tgfb2* (**Figure 4**). However, when cultured with healthy *Sftpc^WT^* fibroblasts, we observed increased expression of AT1 markers in CD44+ and CD44- *Sftpc^C121G^* AT2s, compared to *Sftpc^WT^* AT2s, suggesting a potential role for mesenchymal to epithelial crosstalk in dictating AT2 cell fate in PF.

We acknowledge that alveolar regeneration and maintenance of the alveolar niche is a complex and dynamic process, particularly in the context of PF. While our study was focused specifically on aberrant epithelial-mesenchymal crosstalk that occurs during PF, certainly other cellular compartments including the immune and endothelial cell populations have a critical role in alveolar regeneration following injury (53–57). As such, future studies will incorporate characterization of all lung cellular compartments in the chronic *Sftpc^C121G^* mouse model to compare back to human PF. Further, we acknowledge AT2-to-AT1 differentiation is influenced by several processes, including both biologic factors and mechanical signaling.

In summary, this study demonstrates for the first time that soluble factors from an aberrant alveolar epithelial population that emerges during lung fibrosis can directly promote pathogenic activation of fibroblasts. These findings highlight the utility of the chronic *Sftpc^C121G^* mouse model as an innovative translational tool that recapitulates the clinical course of PF and the emergence of aberrant cell states without the need for non-physiologically relevant extrinsic lung injury. Further, the identification of several profibrotic genes expressed by aberrant transitional alveolar epithelial cells and corresponding fibrotic fibroblast activation, provides a platform in which to test individual or multi-combinatorial therapeutic interventions. While the epithelial paradigm during lung fibrosis has focused on upstream injury and defective progenitor capacity resulting in aberrant lung repair, strategies that modulate the direct epithelial contribution to fibrogenic alveolar niche signaling may provide innovative and effective therapeutics.

## Materials and Methods

### Animals and Mutation Induction

Homozygous *Sftpc^C121G^* mice were achieved through crossing a *Sftpc^C121Gneo^* founder line (Genoway Inc.) with a *Rosa26^ERT2-Cre^* strain, B6.129-Gt(ROSA)26Sortm1(cre/ERT2)Tyj/J (stock no. 008463; The Jackson Laboratory) as previously described (19, 20). Induction of *Sftpc^C121G^* was achieved through oral gavage (OG) tamoxifen (TMX; Millipore Sigma) induction (dissolved in at oil at 20mg/mL) at 200mg/kg in female mice and 230mg/kg in male mice once weekly, as previously described (19). Homozygous *Rosa26^ERT2-Cre^* mice expressing wild-type *Sftpc* (hereafter, *Sftpc^WT^* mice) treated with TMX were used as controls. All mice were weighed once a week during the duration of the study and the TMX dosing was adjusted accordingly. For bleomycin induced injury, lineage tracing AT2 cells was initiated by dosing *Sftpc^CreERT2^*;*Rosa26^tdT^* mice (B6.129S-Sftpctm1(cre/ERT2)Blh/J stock no. 028054; The Jackson Laboratory and B6.Cg-Gt(ROSA)26Sortm14(CAG-tdTomato)Hze/J stock no. 007914; The Jackson Laboratory) with 200mg/kg TMX (in corn oil) by intraperitoneal injections at 14, 12, and 10 days prior to bleomycin administration. Bleomycin was administered at 1.5U/kg intratracheally and mice were monitored daily. mice were housed in an AALAC approved barrier facility at the Perelman School of Medicine (University of Pennsylvania), and under Institutional Animal Care and Use Committee (IACUC) approved protocols (University of Pennsylvania).

### Histology, H&E severity scoring, and immunofluorescence chemistry

Whole lungs were fixed by tracheal instillation of 10% neutral buffered formalin (Sigma) at a constant pressure of 25 cm H_2_O. The Penn Vet Pathology Core Laboratory performed H&E staining on 6μM sections of lung tissues and slides were imaged using an Aperio ScanScope Model: CS2 (Leica) at 40× magnification. Whole lung lobe H&E images were scored for fibrosis severity using a published algorithm (58), as previously described (33, 58, 59). In brief, using the EBImage package in R (60), H&E images were unbiasedly scored for image severity, color coded blue (normal), green (moderate), and severe (red), and each injury score was calculated as the percent of total lung lobe area.

Picrosirius red (PSR) staining of fibrillar collagen was performed on 6μM sections of lung tissues using a PSR Stain Kit (Polysciences). Immunofluorescence chemistry (IFC) staining was performed on 6μM paraffin embedded lung sections using a combination on commercially available primary antibodies and fluorescent AlexaFluor anti-IgG secondary antibodies (**Supplementary Table 1**). Images were obtained using an Eclipse Ti2 Series inverted microscope (Nikon). Images were analyzed using Fiji 2.15.1.

### Single-cell RNA sequencing and analysis

Single cell suspensions from end-stage (Week 7) *Sftpc^C121G^* or *Sftpc^WT^* were obtained as previously described (20, 33). Perfused whole lungs were diced using surgical scissors and digested in a Phosphate Buffered Saline (PBS; Mg^-^Ca^-^) solution supplemented with 5mg/mL Collagenase Type I (Thermo) and 50 units of DNase (Millipore Sigma) at 37°C for 1 hour with frequent mechanical perturbation. Single cell suspensions were subsequent passed through a 70 μm nylon mesh filter and red blood cells were removed using an ACK Lysis Buffer (Thermo). To capture a cell suspension containing an equal mixture of strictly epithelial and mesenchymal cells, each lung was divided into two samples and coated with either CD45, CD31, and EpCAM antibody conjugated microbeads (Miltenyi Biotec) to negatively select for mesenchymal cells or EpCAM antibody conjugated microbeads to positively select for epithelial cells. Microbead-coated cells were then passed through magnetic LS columns (Miltenyi Biotec), and both mesenchymal and epithelial cell populations from each lung were pooled back together at a 50:50 ratio before loaded onto individual GemCode instrument (10× Genomics; 2 for each genotype). Single-cell barcoded droplets were produced using 10X Single Cell 3′ v3 chemistry. Libraries generated were sequenced using the HiSeq Rapid SBS kit, and the resulting libraries were sequenced across the 2 lanes of an Illumina HiSeq2500 instrument in a high-output mode. scRNA-Seq reads were aligned to mouse genome (mm10/GRCm38) using STARSolo (version 2.7.5b). After initial quality control and processing, we analyzed the scRNA-Seq data using the Scanpy pipeline (61). Genes expressed in fewer than 3 cells were removed, and cells with fewer than 200 genes and a mitochondrial fraction of less than 20% were excluded. Counts were log normalized using scanpy.pp.normalize_per_cell (counts_per_cell_after=1×104), followed by scanpy.pp.log1p. To integrate data from multiple samples, we used Scvi-tools (62). We applied scvi.model.SCVI. setup_anndata() to establish the model parameters for integration, including: layer, categorical_covari-ate_keys and continuous_covariate_keys. We then performed a principal component analysis (PCA) and generated a K-nearest neighbor (KNN) graph using scanpy.pp.neighbors with n_neighbors=15. The resulting KNN graph was used to perform Uniform Manifold Approximation and Projection (UMAP) dimension reduction to visualize the cells in 2 dimensions using scanpy.tl.umap(). Clustering was per-formed using the Leiden algorithm with scanpy.tl.leiden (63). We identified cell populations using known canonical marker genes (**Supplemental Table 2**) or by assessing cluster-defining genes based on differential expressions. Trajectory analysis was performed on the UMAP reduction using the R Slingshot package with both AT2 cells and the UPR Active AT2 cells as a starting point without any assigned end points. The lineages are visualized using a custom function in R. Analysis of ligand-receptor signaling was performed using NICHES v1.0.0 as previously described(51). Seurat v4.3.0, ggplot2, and ComplexHeatmap v2.14.0 were used to plot selected significant findings, trends, and patterns of interest.

### Fluorescence-activated cell sorting (FACS) of epithelial and mesenchymal populations

Single cell suspensions of blood-free whole lung cells were obtained as described above. Fluorophore-conjugated antibodies were used to stain lung cell populations (**Supplementary Table 1)**. All cells were sorted using a CytoFlex SRT (Beckman) and cells were captured in ice-cold FACS buffer (0.1% BSA, 2mM EDTA, and PBS pH 7.4). *Sftpc^WT^*AT2s and *Sftpc^C121G^* AT2/transitional cells were defined as (CD45^-^;EpCAM^+^;CD104^-^;CD200^+^). *Sftpc^WT^*AT2s and *Sftpc^C121G^* bulk fibroblasts were isolated as (CD45^-^;EpCAM^-^;CD31^-^;MCAM^-^;Pdgfra^+^). Fluorescent IgG isotype controls were used to determine staining positivity. Flow cytometry gating and quantification was performed using FlowJo v10.10 software.

### RNA isolation, bulk RNA sequencing, and RT-PCR

RNA was isolated from freshly sorted mouse lung cells, or cells derived from cell culture assays, using an RNeasy Mini Kit (Qiagen) following the manufacturer’s protocol and RNA quantification was performed using a NanoDrop One (Thermo Fisher Scientific). RNA was either processed for bulk RNA sequencing or reverse transcribed using a Verso cDNA Synthesis Kit (Thermo Fisher Scientific). For RNA sequencing, library preps were performed by Penn NextGen Sequencing Core. Fastq files were evaluated for quality control with the FastQC program and then aligned against the mouse reference genome (mm10) using the STAR aligner87. Duplicate reads were flagged with the MarkDuplicates program from Picard tools and excluded from analysis. Per gene read counts for Ensembl (v67) gene annotations were computed using the R package Rsubread. Gene counts, represented as counts per million (CPM), were nominalized using TMM method in the edgeR R package, and genes with 25% of samples with a CPM < 1 were considered low expressed and removed. The data were transformed with the VOOM function from the limma R package to generate a linear model and perform differential gene expression analysis. We employed the empirical Bayes procedure as implemented in limma to adjust the linear fit and to calculate P values given the small sample size of the experiment. We adjusted P values for multiple comparisons using the Benjamini-Hochberg procedure. Heatmaps were generated using Morpheus (https://software.broadinstitute.org/morpheus). Protein-protein interactions were obtained using STRING (Search Tool for Retrieval of Interacting Genes/Proteins) database. Gene ontogeny analysis was performed using the Database for Annotation, Visualization, and Integrated Discovery (https://david.ncifcrf.gov/home.jsp) based on differentially enriched genes (FC >1.5 P-value <0.05). Key pathway analyses was performed on gene lists identified from the GSEA molecular signature database.

### Primary Mouse Organoid Culture and Analysis

Freshly sorted alveolar epithelial or mesenchymal cell populations were isolated from whole lungs and sorted by FACS as described above. Mouse lung organoids were formed according to previously established protocols (64). Various combinations and *Sftpc^WT^* AT2s and CD44+/- *Sftpc^C121G^* AT2/transitional cells and Pdgfra^+^ fibroblasts were utilized where states. Regardless of cell origin, each technical replicate contained 5000 alveolar epithelial cells and 50,000 Pdgfra^+^ fibroblasts. Cell suspensions were resuspended in a mix of 50% GFR-Matrigel (Corning) and 50% Small Airway Epithelial Cell Growth Media (SAGM; Lonza) and plated in a 24-well Falcon Cell Culture Insert (Falcon). The cell/Matrigel culture solidified for 30 minutes at 37°C before 500 µl SAGM was added to the basal layer of the transwell plate. SAGM was made fresh for each experiment in accordance with the manufacturer’s instructions, using a Lonza BulletKit (with the exception of excluding Hydrocortisone, BSA, Triiodothyronine, and Epinephrine). For the first two days of culture SAGM was additionally supplemented with 10 μM rock inhibitor (Y-27632 dihydrochloride, Sigma). Subsequently, SAGM was refreshed every other day for the duration of the experiment.

AT2 alone organoid culture was performed as previously described (46, 47). Freshly flow sorted AT2 cells were resuspended at GFR-Matrigel at 100 cells/μL and 50 μL Matrigel droplets were plated in a 12-well plate (Falcon). Matrigel droplets were allowed to solidify for 30 minutes at 37°C. For media, previously published AT2 Maintenance Media (AMM) (46, 47), with minor adjustments, was utilized. AMM was composed of 10 μM SB431542 (Abcam), 3 μM CHIR99021 (Tocris), 1x Insulin-Transferrin-Selenium (ITS) (Thermo), 15mM HEPES (Thermo), 1x Glutamax (Thermo), 50ng/mL hEGF (Gibco), 5μg/mL Heparin (Sigma), 1x B27 supplement (Gibco), 1x N2 supplement (Gibco), 1.25mM N-Acetyl Cysteine (Sigma), and varying concentrations of FGF7 (Tocris) as described, in Advanced DMEM/F12 (Gibco). For the first two days of culture AMM was additionally supplemented with 10 μM rock inhibitor (Y-27632 dihydrochloride, Sigma). Subsequently, AMM was refreshed every other day for the duration of the experiment.

All organoid imaging was performed using an inverted Leica Thunder microscope and images were quantified for organoid size and colony forming efficiency (CFE) using Fiji 2.15.1.

### EpCAM+ Magnetic Bead Enrichment from Organoid Cultures

To enrich epithelial and mesenchymal cells out of organoid culture for gene expression analysis, Matrigel droplets containing organoids were initially dissociated in 2mg/mL Dispase (Gibco) for 30 minutes at 37°C. Isolated organoids were subsequently resuspended in Accutase (Simga) for 15 minutes at 37°C to receive a single cell suspension. Single cell suspensions were resuspended in FACS Buffer with biotinylated anti-mouse CD326 (EpCAM; Biolegend) for 30 minutes at 4°C. Excess antibody was washed off in FACS Buffer and cell pellet was resuspended in FACS Buffer containing Dynabeads M-280 Streptavidin (Intvitrogen) for 30 minutes at 4°C. Tubes containing cell and bead suspensions were then placed on a MagJet Rack (Thermo) to separate EpCAM+ (on-bead) and EpCAM- (in suspension) fractions. EpCAM-suspensions were pelleted before RNA isolation as described above. EpCAM+ (on-bead) fractions were lysed in RLT Buffer (Qiagen) and centrifuged at 300xg for 5 minutes to pellet beads. Lysed supernatant was then processed using an RNeasy Mini Kit (Qiagen), according to the manufacturers protocol.

### Primary Mouse 2D Culture

Alveolar epithelial cells sorted by FACS as described above or heathy alveolar fibroblasts from *Sftpc^WT^* mice were sorted as (Pdgfra^+^;Sca1^-^;CD45^-^;CD31^-^:EpCAM^-^;MCAM^-^) and plated on 24-well Falcon Cell Culture Inserts (Falcon) that had previously been coated with hESC-qualified Matrigel at room temperature for 2 hours. The apical compartment of each transwell was loaded with 500 µl DMEM:F12 (10% FBS, Glutamax, and Primocin) containing 1.0 × 10^5^ alveolar epithelial cells or 2.0 × 10^5^ alveolar fibroblasts. The basal compartment was additionally loaded with 500 µl of the same media. For alveolar epithelial cells, media supernatant was collected and cell monolayers were lysed for RNA isolation at Day 5. Alveolar fibroblasts were cultured for 2 days before media was changed to conditioned media derived from alveolar epithelial cell cultures. At 2 days post-supernatant exposure, alveolar fibroblasts were lysed for RNA isolation.

### Statistics

All data are presented with dot plots and group mean ± SEM, unless otherwise indicated. Statistical analyses were performed with GraphPad Prism. Student’s t test (1 or 2 tailed, as appropriate) were used for 2 groups. Multiple comparisons were done by 1-way ANOVA, which was performed with post hoc testing as indicated; survival analyses was performed using Kaplan-Meier with Mantel Cox correction. In all cases, statistical significance was considered at P ≤ 0.05.

### Data and code availability

The sequencing data generated in this study are deposited in Gene Expression Omnibus (GEO) and will be made available upon publication of this work.

## Supplemental Figures

**Supplemental Figure 1.**
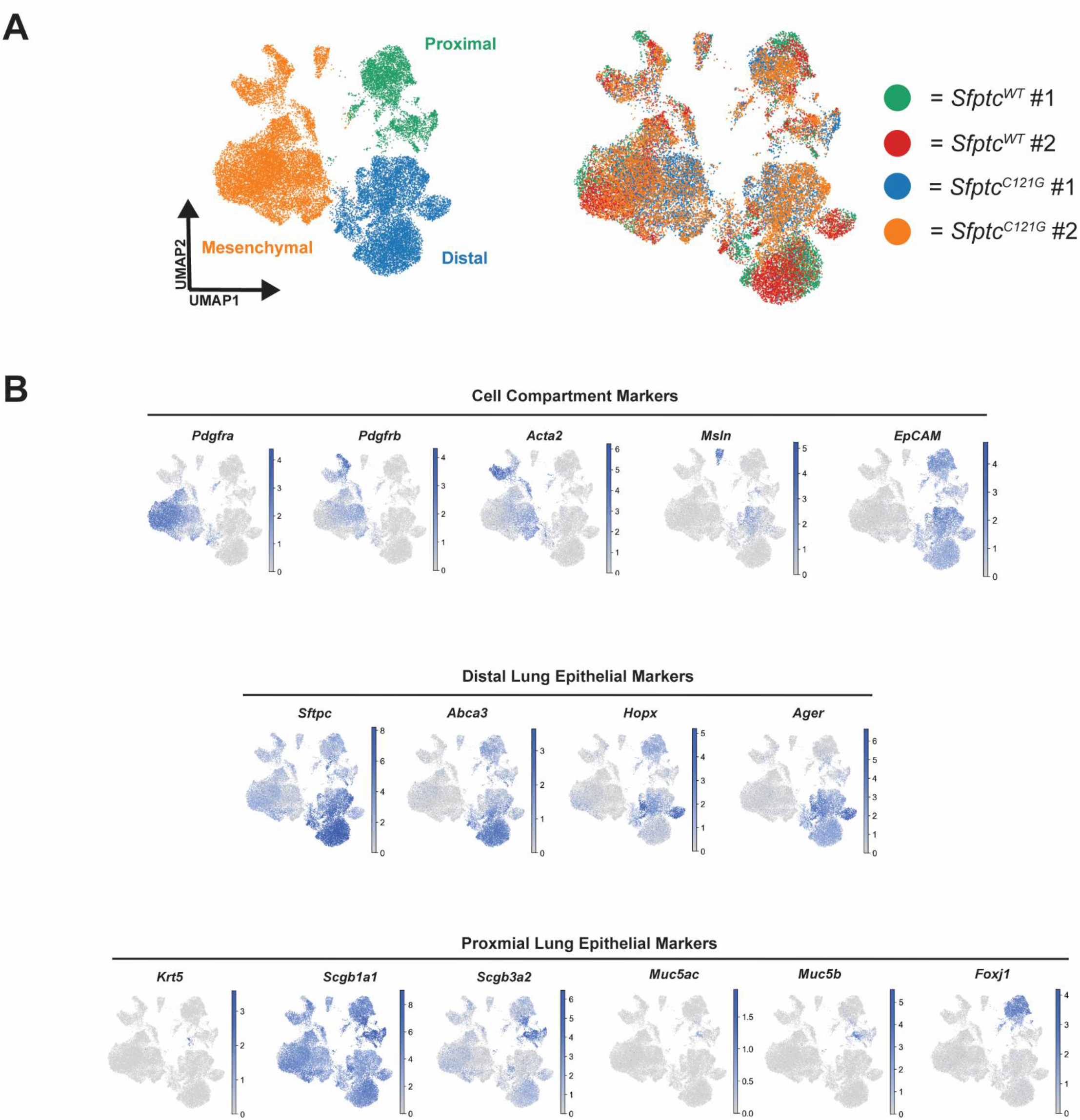
All samples contributed to clusters of mesenchymal and epithelial cell compartments. (A) Representative UMAP showing distribution of individual sample clustering. (B) Gene expression UMAPs of lineage-specific marker genes on total cell populations.

**Supplemental Figure 2.**
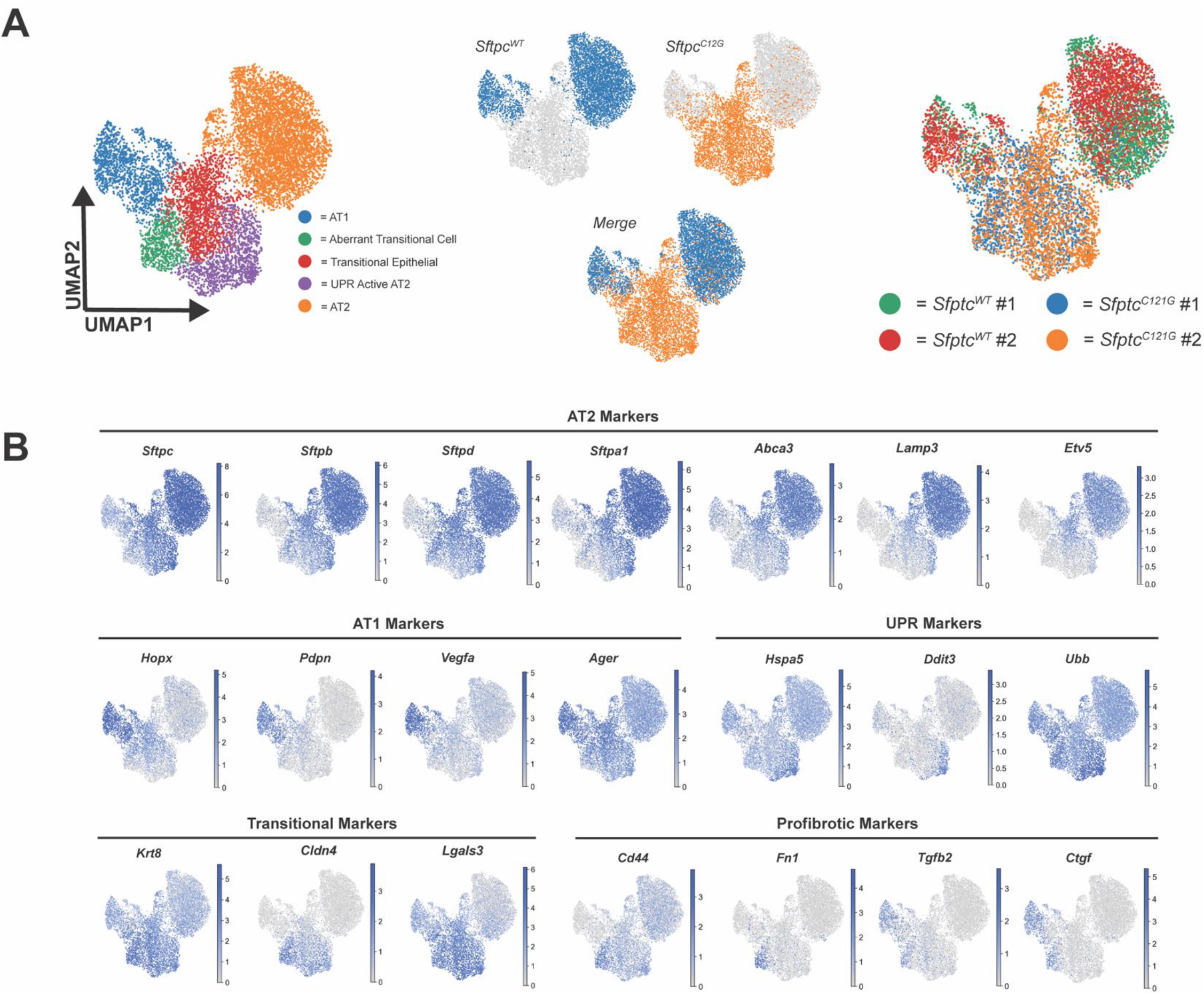
Re-clustered distal lung epithelium demonstrated distinct epithelial population in *Sftpc^WT^* and *Sftpc^C121G^* mice. (A) Representative UMAP showing distribution of individual condition and sample clustering. (B) Gene expression UMAPs of lineage-specific marker genes on distal epithelium populations.

**Supplemental Figure 3.**
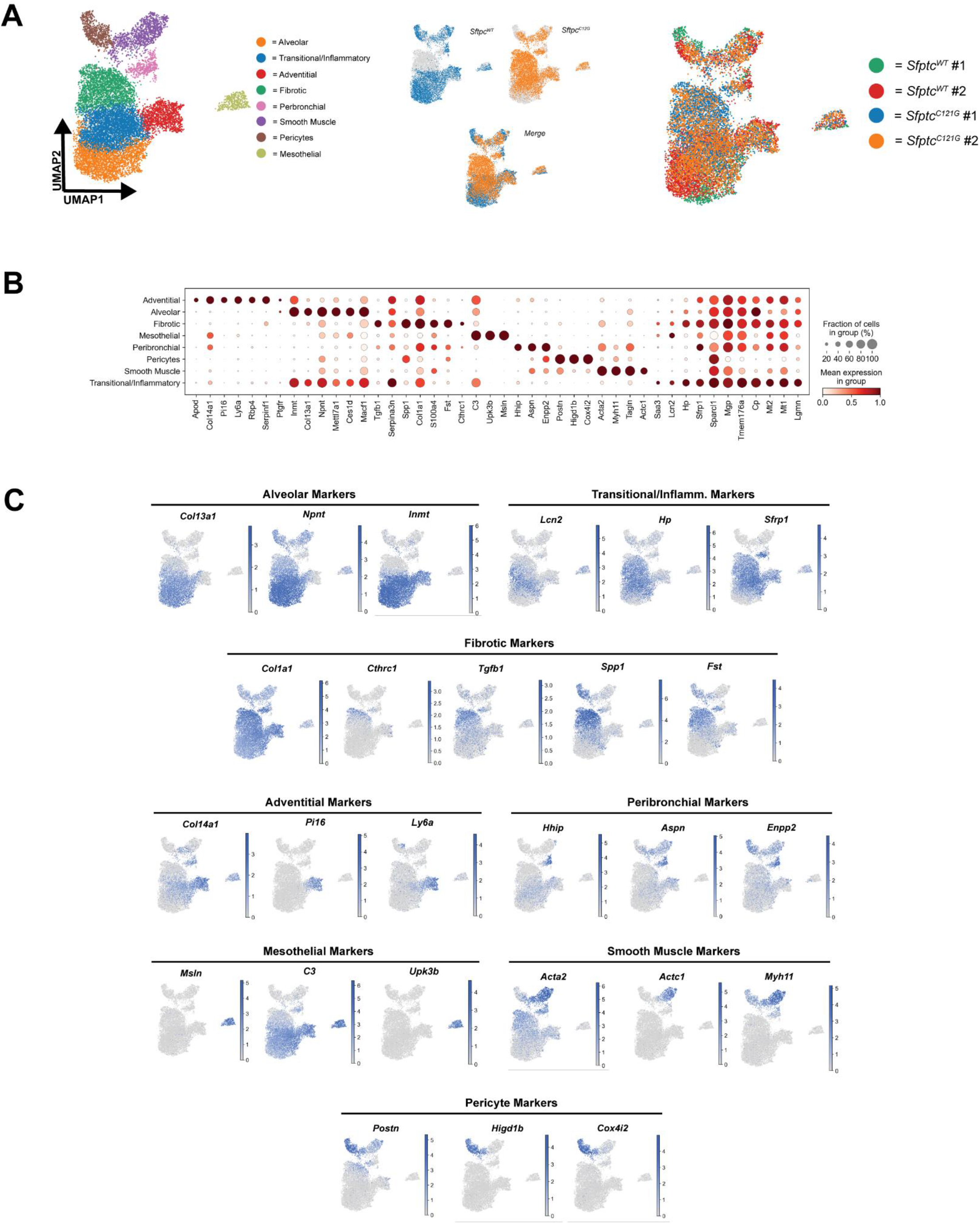
Re-clustered mesenchymal populations identify recognized fibrotic populations in *Sftpc^C121G^* mice. (A) Representative UMAP showing distribution of individual sample clustering. (B) Dot plot exhibited lineage-specific markers gene amongst mesenchymal populations. (C) Gene expression UMAPs of lineage-specific marker genes on mesenchymal populations.

**Supplemental Figure 4.**
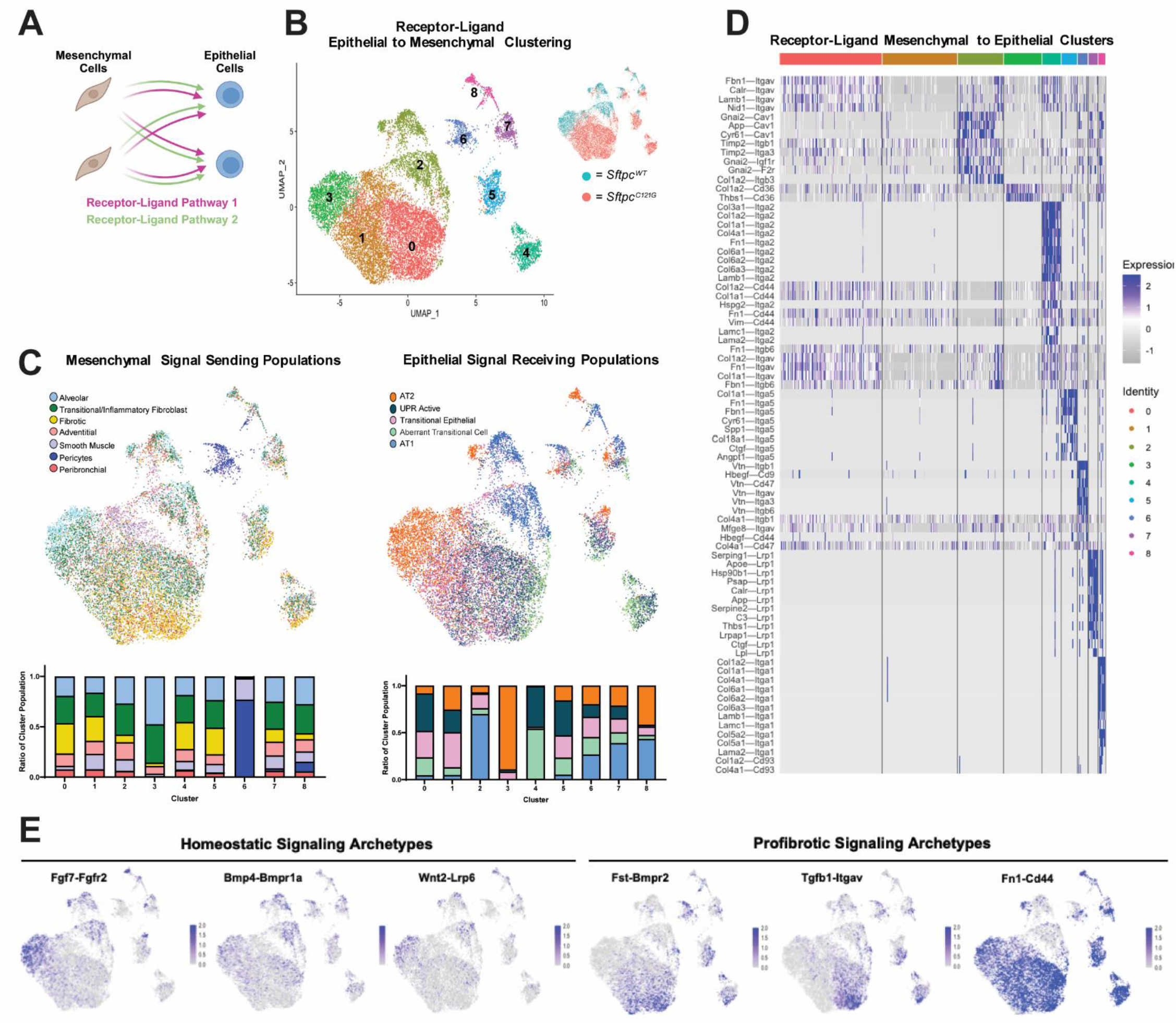
Mesenchymal-to-epithelial signaling is characterized by a loss of homeostatic signaling and gain of profibrotic signaling. (A) Schematic demonstrating single-cell ligand-receptor mapping using NICHES. (B) UMAP representation of unbiased mesenchymal-to-epithelial ligand-receptor mapping clustered dependent on distinct signaling archetypes. (C) UMAPs labeled by mesenchymal sending cell type and epithelial receiving cell type populations. Ratios of each cell type within a given cluster is provided. (D) Heatmap of individual ligand-receptor pairs that define each cluster archetype. (E) Representative UMAPs of known homeostatic and profibrotic ligand-receptor pairs that are altered in fibrosis.

**Supplemental Figure 5.**
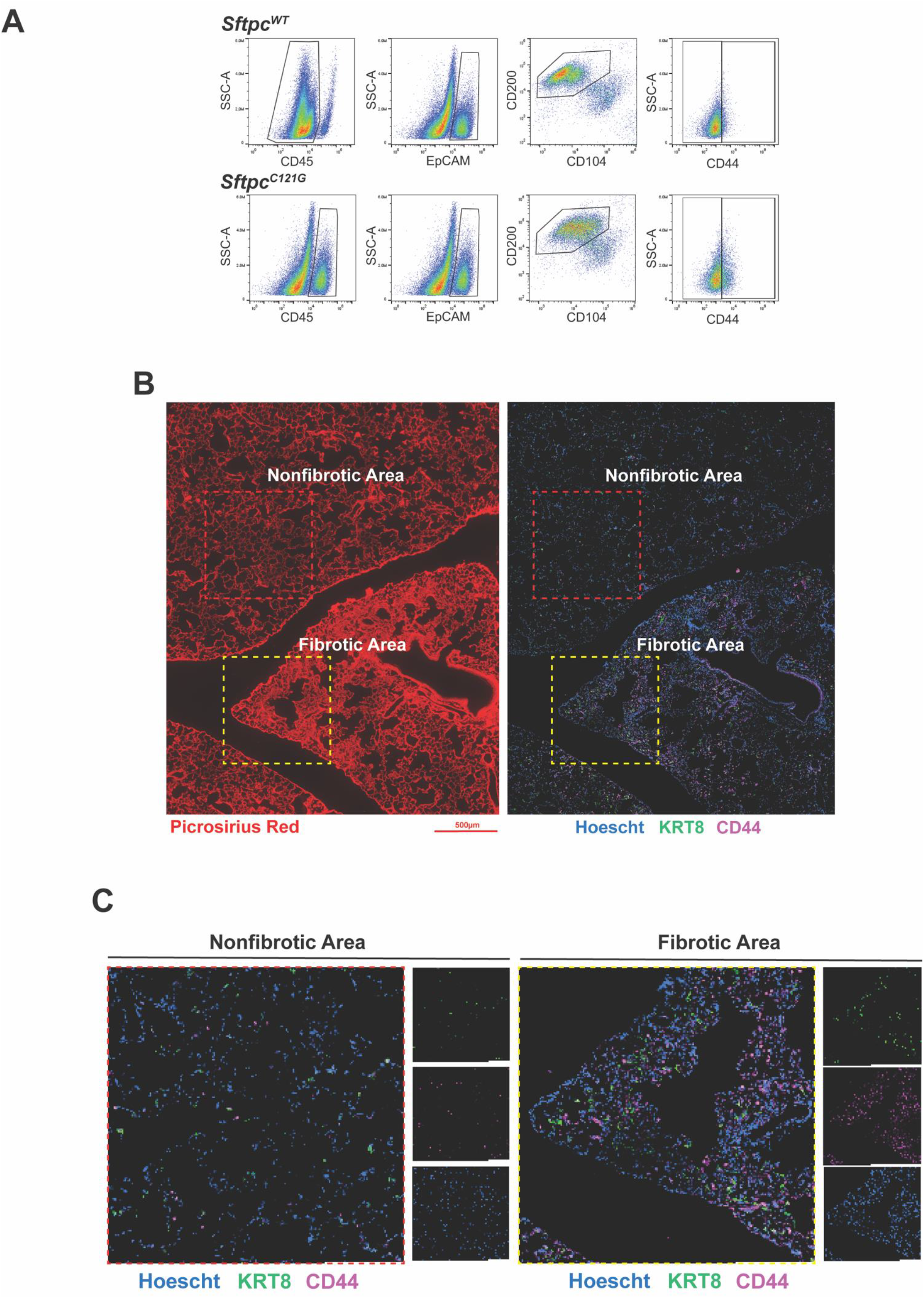
CD44+ transitional cells are present in fibrotic areas of *Sftpc^C121G^* lungs. (A) Representative flow cytometry gating for isolating CD44+/- AT2/transitional cells in *Sftpc^WT^* and *Sftpc^C121G^* lungs (B) PSR staining and Hoescht, KRT9, and CD44 staining in *Sftpc^C121G^* lungs showing CD44+/KRT8+ cells are present in fibrotic foci. Scale bar = 500 µm. (B) Insets from non-fibrotic and fibrotic areas of *Sftpc^C121G^* lungs.

**Supplemental Figure 6.**
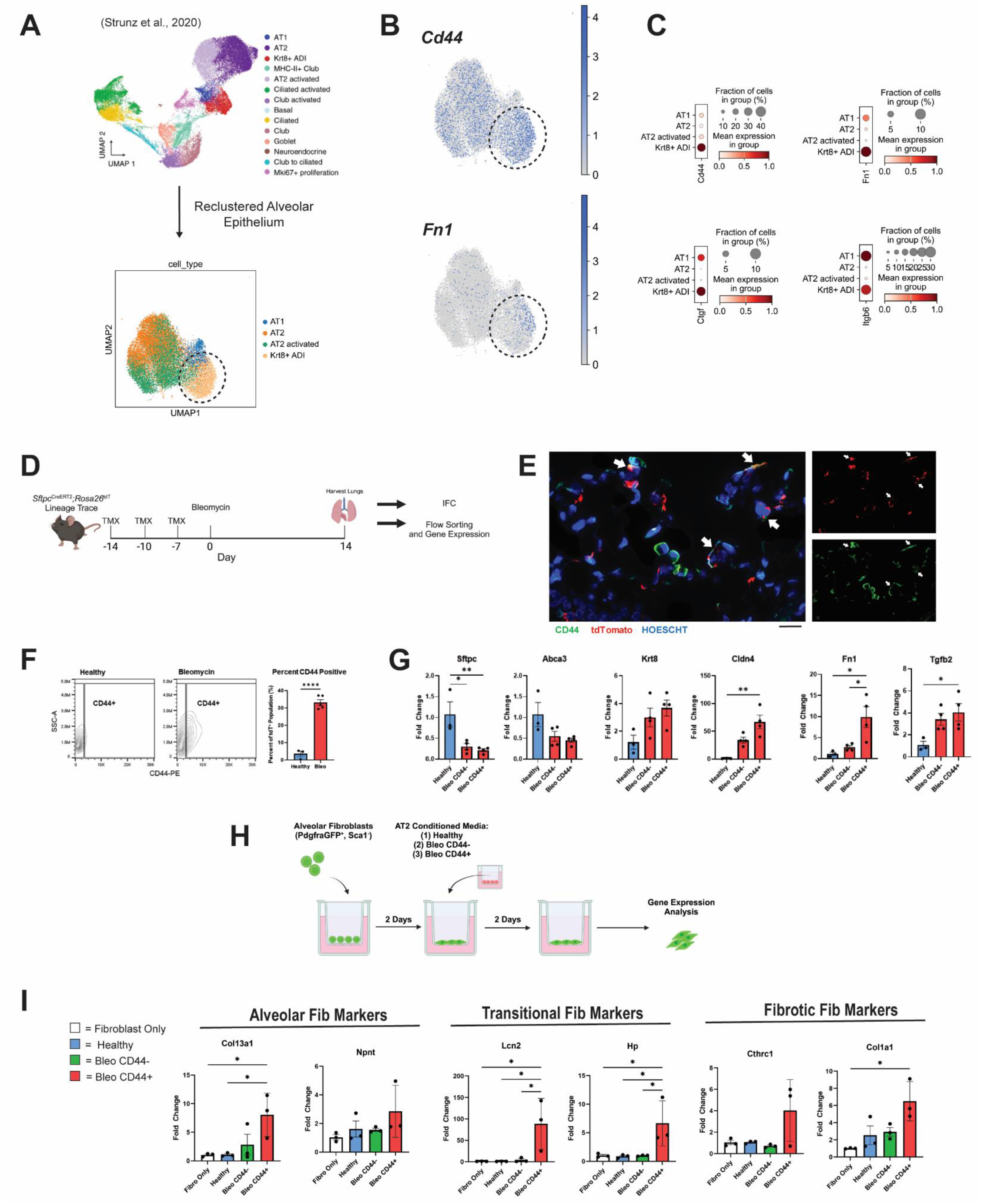
*Sftpc* lineage traced CD44^+^ cells in a bleomycin model recapitulate profibrotic gene expression. (A) Previously published UMAP representation of epithelial cell types captured by scRNAseq at 18 time points (Strunz et al., 2020), and re-clustering of the alveolar epithelium. (B) UMAP representation and (C) Dot plot gene scores according to alveolar epithelium cell clusters. (D) Schematic tamoxifen (TMX) dosing and bleomycin administration in *Sftpc* lineage traced mice prior to Day 14 analysis. (E) IFC demonstrating tdT^+^ CD44+ elongated cells at Day 14 in bleomycin model. Scale bar = 20 µm (F) Flow cytometry analysis of CD44 upregulation in mice that received bleomycin at Day 14. Student’s t-test, **** P<0.0001. (G) Gene expression analysis of total healthy tdT^+^ AT2s compared to CD44^-^ and CD44^+^ tdT^+^ cells at Day 14 in mice that received bleomycin. (H) Schematic of healthy alveolar fibroblast exposure to conditioned supernatant from AT2s 2D cultured for five days. (I) Gene expression of alveolar fibroblasts after two days of supernatant exposure. Statistical analysis by one-way ANOVA (*p < 0.05).

**Supplemental Figure 7.**
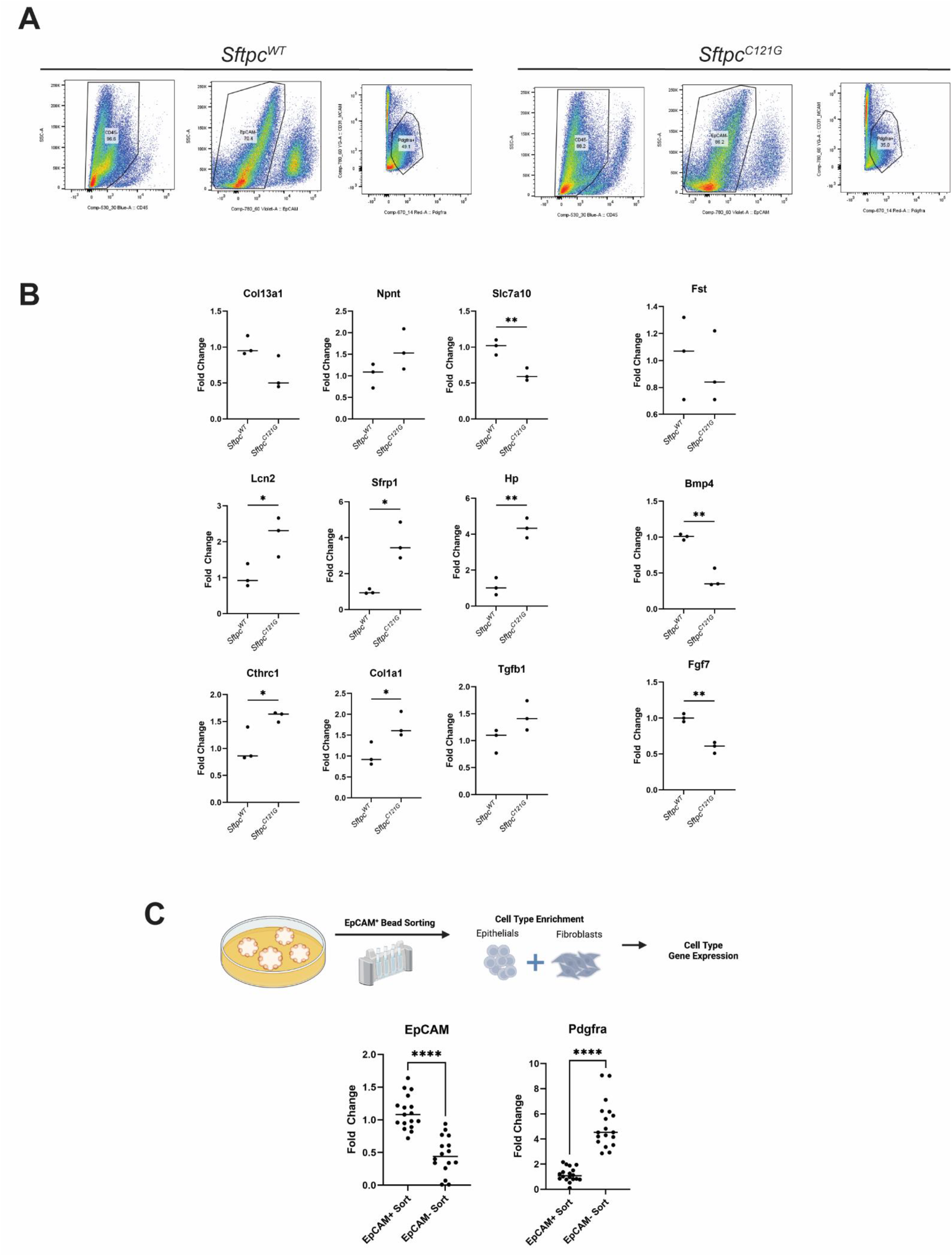
Freshly sorted bulk Pdgfra^+^ fibroblasts from *Sftpc^C121G^* mice have increased expression of transitional and fibrotic gene markers. (A) Representative flow cytometry gating of Pdgfra^+^ fibroblasts from *Sftpc^WT^* and *Sftpc^C121G^* mice. (B) Gene expression analysis of fibroblast gene markers. (C) Schematic of positive EpCAM-selection from mixed culture organoid with gene expression analysis from EpCAM^+^ and EpCAM^-^ populations. Statistical analysis by student’s t-test (*p < 0.05, ** p<0.005, **** p<0.0001).

**Supplemental Table 1.**
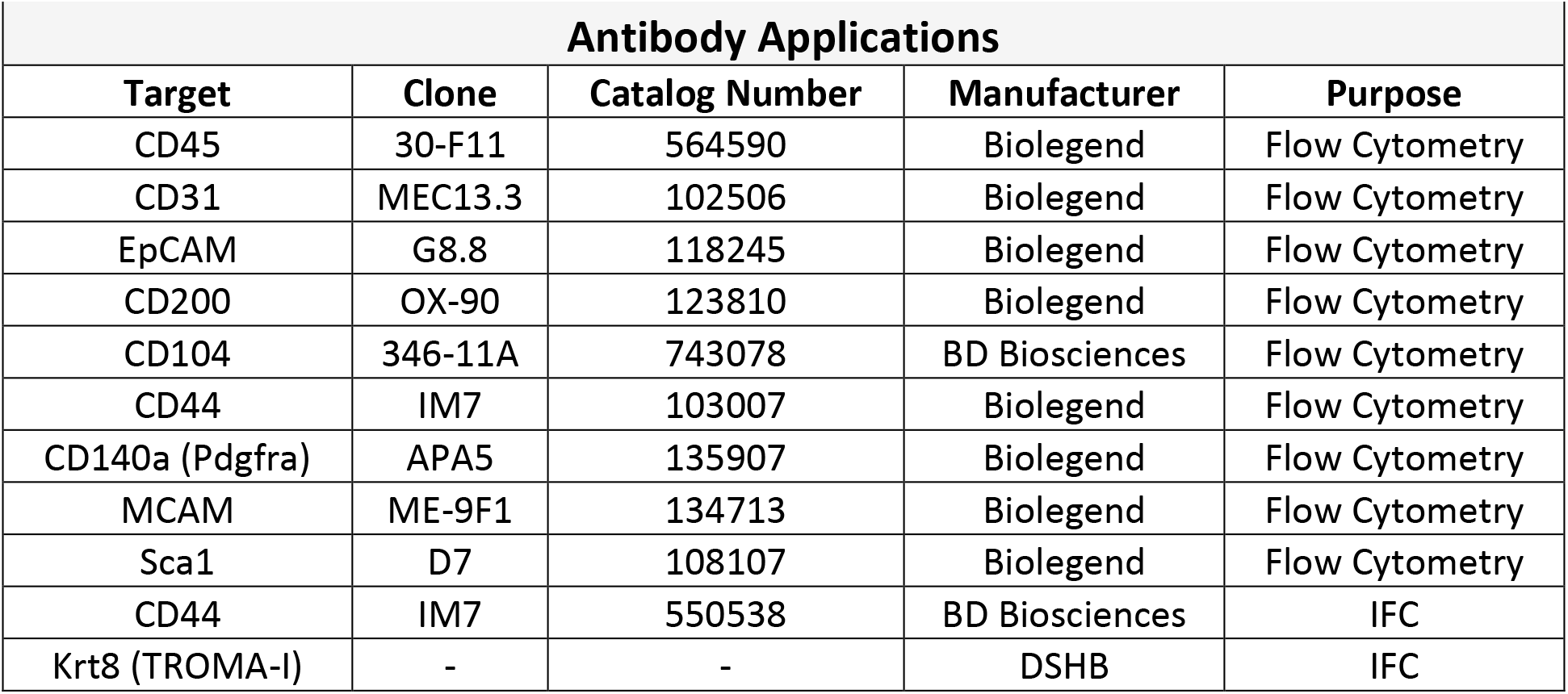

**Supplemental Table 2.**
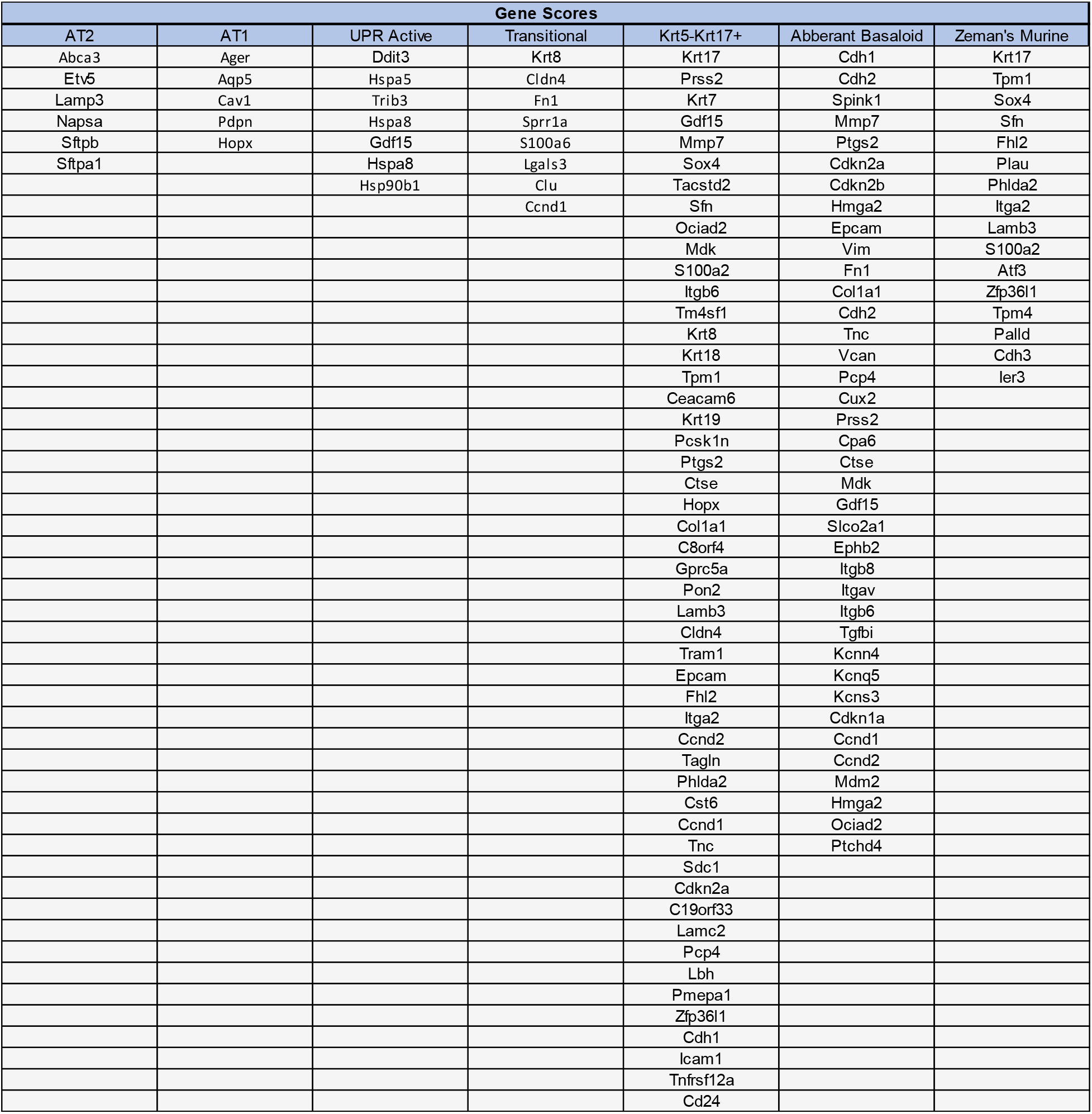

